# Hexosyl transfer *via* a covalent intermediate within a dual retaining and inverting enzyme

**DOI:** 10.64898/2026.10.20.760405

**Authors:** Mert Sağıroğlugil, Beatriz Piniello, Miron A. Leanca, Eduardo S. Kitano, Courtney J. Mycroft-West, Georgios Papadakis, Maria Alexandra Gafitescu, Shabaz Mohammed, Benjamin G. Davis, Carme Rovira, Liang Wu

**Affiliations:** Departament de Química Inorgànica i Orgànica (Secció de Química Orgànica) and Institut de Química Teòrica i Computacional (IQTCUB), Universitat de Barcelona, Martí i Franquès 1, 08028 Barcelona, Spain; The Rosalind Franklin Institute, Harwell Science C Innovation Campus, OX11 0QX, Didcot, UK; Department of Biochemistry, University of Oxford, OX1 3QU, Oxford, UK; Department of Chemistry, University of Oxford, OX1 3TA, Oxford, UK; Department of Pharmacology, University of Oxford, OX1 3QT, Oxford, UK; Institució Catalana de Recerca i Estudis Avançats (ICREA), Passeig Lluís Companys 23, 08020 Barcelona, Spain

## Abstract

Bifunctional glycosyltransferases underpin biosynthesis of essential glycans, including bacterial capsular antigens, and eukaryotic glycosaminoglycans. Typically, these enzymes transfer hexose sugars from activated nucleotide-phosphate donors, forming new glycosidic bonds with either retention or inversion of anomeric stereochemistry. Retaining hexosyl transfer typically proceeds *via* non-covalent *S_N_i*-type pathways, but recent studies of octulosyl Kdo-transferases have reignited debate over covalent mechanisms. Here we comprehensively dissected the activity of the archetypal bifunctional glycosyltransferase PmHS2, providing X-ray structures of ligand complexes alongside atomistic simulations to delineate full substrate-to-product reaction trajectories. Combined computational, biochemical and evolutionary evidence suggests that PmHS2 retaining GlcNAc-transfer likely proceeds *via* a previously undescribed covalent mechanism, involving considerable assistance from departing nucleotide-phosphate rotation. In mapping reaction pathways, we also report the first example of a glycosyl donor substrate coopted as an acceptor, which enables enzymatic self-priming for polymerization. These new covalent and self-priming functionalities highlight unexplored routes to modulate or inhibit polysaccharide biosynthesis.

## Introduction

Precise enzymatic processing of carbohydrates is a vital biological process, which underpins their myriad structural and signaling functions across all domains of life^1–4^. The central importance of carbohydrates in biology is underscored by the substantial metabolic machinery dedicated towards their regulation^5^: ∼1–5% of genes in any organism encode for carbohydrate active enzymes, which together coordinate chemo-, regio- and stereoselective transformations to produce complex glycan structures^6^. Resolving the catalytic mechanisms of carbohydrate active enzymes is central to understanding their biological functions, as well as that of their glycan products, in turn opening the door for the modulation of both.

Glycosyltransferases (GT) are the principal enzymes responsible for generating new glycosidic bonds, which catalyze the transfer of sugars from (typically) phosphate-linked donors onto defined acceptor substrates. Mechanistically, inverting GTs that transfer sugars with inversion of anomeric configuration are understood to utilize single-step S_N_2-type mechanisms, in which a basic enzyme residue (usually Asp/Glu) assists glycosyl acceptor attack upon a donor anomeric center, from the opposite side to its activated leaving group (**Figure 1a**)^7,8^. By contrast, retaining GTs principally operate *via* S_N_i-type internal return, wherein a constrained acceptor approaches the donor from the same ‘face’ as its leaving group, and so is aided by concomitant deprotonation by the departing donor phosphate (**Figure 1b**)^9,10^.

**Figure 1.**
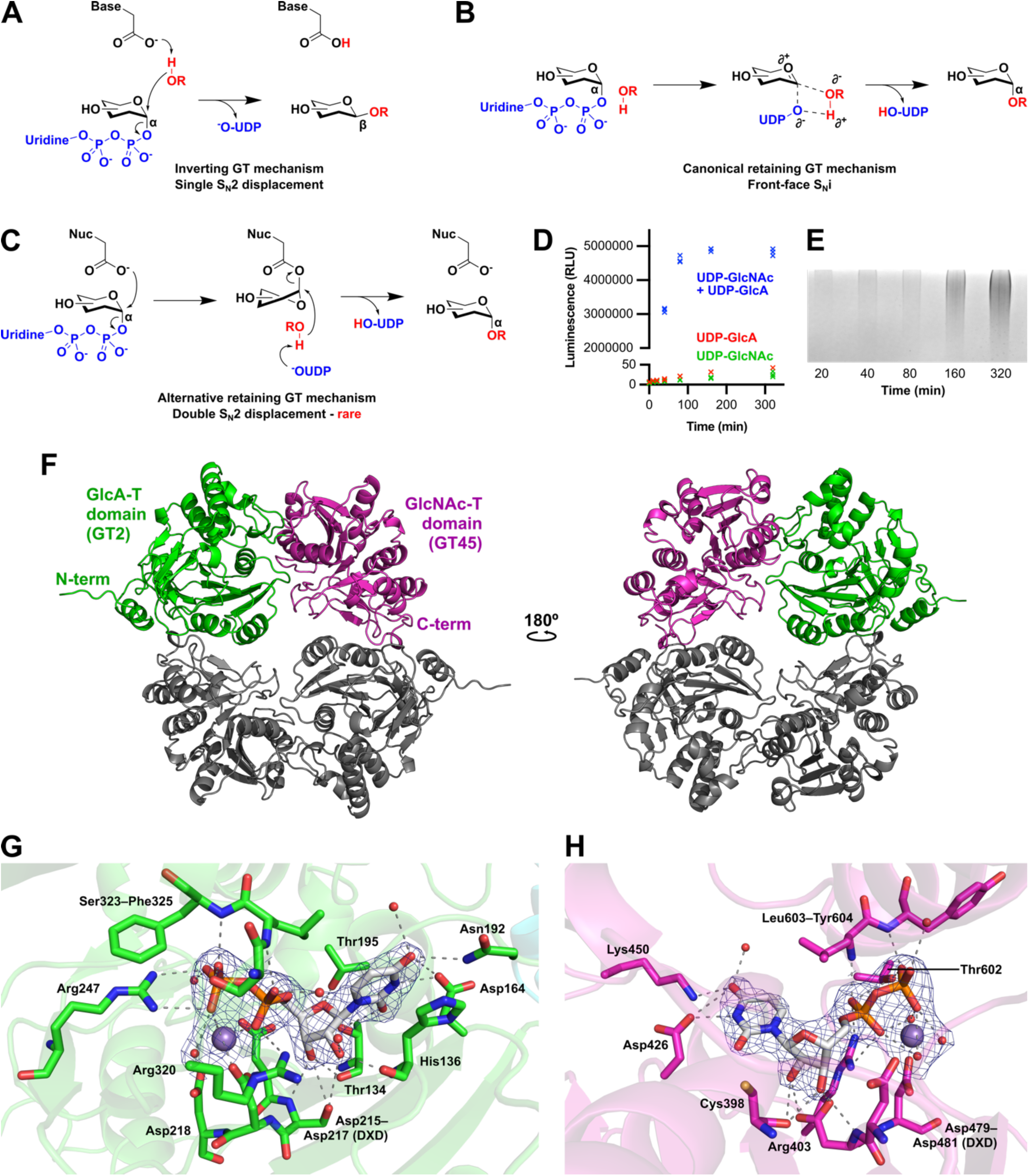
Overview of Δ80-PmHS2 structure and function. Schematic of canonical inverting (**A**) and retaining (**B**) GT mechanisms, involving bond formation by single S_N_2 displacement and S_N_i-like displacement, respectively. (**C**) Proposed double S_N_2 displacement mechanism for retaining GTs, with transient formation of a covalent enzyme adduct. (**D**) UDP-Glo assay measuring Δ80-PmHS2 activity in the presence of one or both UDP-sugar donors. (**E**) Alcian blue stained SDS-PAGE gel^27^, showing production of heparosan over time by Δ80-PmHS2, co-incubated with UDP-GlcNAc and UDP-GlcA. (**F**) Ribbon representation of the Δ80-PmHS2 homodimer. GlcA-T and GlcNAc-T domains of one protomer are depicted in green and pink respectively. The homodimer partner is depicted in grey. (**G**, **H**) UDP interactions within the PmHS2 GlcA-T and GlcNAc-T domains. Waters -red spheres. Mn^2+^ - grey spheres. Electron densities are REFMAC σ_A_-weighted 2mFo–DFc, contoured to 1.5 σ (0.2C e^−^.Å^−3^).

With analogy to glycoside hydrolases that cleave sugar bonds^11^, glycosyltransfer mechanisms involving covalent intermediates have been proposed for retaining GTs carrying well-placed nucleophilic residues within their active sites. Such ‘double displacement’ reactions may be further aided by the formation of stabilized oxocarbenium-like glycosyl intermediates, which support migration of the sugar to/from a nearby enzyme-contributed nucleophile (**Figure 1c**). Recently, bacterial eight-carbon sugar (octulose) Kdo-transferases that employ covalent mechanisms have been reported^12,13^, which likely exploit the intrinsically stabilized Kdo intermediate that arises from neutralization of its oxocarbenium-like cation by an adjacent C1 carboxylate. In contrast, six-carbon (hexose) sugars that are most widespread across biology typically lack such charge-neutralization mechanisms, thus GTs that processes these substrates must utilize higher-energy pathways, with correspondingly reduced stability and lifetime. To date, only a few retaining galactosyltransferases and N-acetylgalacosyltransferases^14,15^ have been proposed to operate *via* covalency, with long-standing debate surrounding each example^16,17^.

Many essential biological carbohydrates are comprised of repeating disaccharide units, including bacterial capsular polysaccharides, and eukaryotic glycosaminoglycans such as heparan and chondroitin sulfate. Such polymers are often constructed by bifunctional GTs that can transfer both saccharides; these likely represent catalytically optimized systems, wherein each glycosyl transfer step must be coordinated with its functional neighbor. One such bifunctional GT, PmHS2 from *Pasteurella multocida*, is an archetypal heparosan ([GlcNAc-GlcA]_n_) synthase that harbors both inverting glucuronic acid (GlcA)-transferase (GlcA-T; CAZy family^6^ GT2) and retaining N-acetylglucosamine (GlcNAc)-transferase (GlcNAc-T; CAZy family GT45) activities within the same protein chain^18,19^. Owing to its widespread use in chemoenzymatic heparosan and heparan sulfate production^20,21^, PmHS2 has been the subject of extensive functional studies^22^, with recent cryo-EM and X-ray structures revealing a compact dimer structure carrying extensively separated GlcNAc-T and GlcA-T catalytic domains^19^ reminiscent of eukaryotic bifunctional GTs such as EXT1/EXT2^23,24^, CHSY3-CHPF^25^, and EXTL3^26^. Notably, these structural studies of PmHS2 revealed a poised nucleophilic Asp569 within its retaining GlcNAc-T domain, raising, for us, the possibility of a rare covalent mechanism^18,22^. However, despite these suggestions of potential mechanistic novelty, molecular understanding of PmHS2 (and other bifunctional GT) activities has remained limited, owing to a broader lack of structural insights that can inform upon enzyme/substrate interactions.

Here, we present a comprehensive dissection of the PmHS2 catalytic cycle. By capturing a complete set of PmHS2 ligand complexes across both of its catalytic domains, we have mapped detailed substrate-to-product trajectories for both its GlcA and GlcNAc transfer reactions, resolving ligand interactions of key importance for catalysis. Combining experimental structures with molecular dynamics (MD) and quantum mechanics/molecular mechanics (QM/MM) simulations, we further analyze the PmHS2 retaining GlcNAc-T reaction in atomistic detail. Our results not only suggest that PmHS2-catalyzed GlcNAc-transfer proceeds *via* key covalent intermediates, but also that substrate plasticity in its GlcA-T domain enables functions such as self-priming of polymerization. Collectively, our data highlight the considerable undiscovered diversity underpinning GT mechanisms in nature.

## Results

### Ligandable crystal form of a functional PmHS2 homodimer

To examine PmHS2 inverting and retaining catalytic mechanisms, we expressed and purified a known stabilized and highly activated enzyme variant lacking its initial 80 amino acids (Δ80-PmHS2)^22^. Catalytic function of this construct was examined by incubation with UDP-GlcA and UDP-GlcNAc sugar donors, confirming optimal heparosan biosynthesis in the presence of both substrates (**Figure 1d, e**). SEC-MALS identified a predominant ∼128 kDa homodimeric species for Δ80-PmHS2 in solution (monomer ∼67 kDa), with a smaller amount of tetramer (∼237 kDa) at a ratio of ∼4:1 dimer:tetramer (**Supplemental Figure 1**).

Limited insights into PmHS2 substrate or product interactions to date reflect the need for new structural forms, which are more amenable to liganding. We found that Δ80-PmHS2 pre-complexed to Mn^2+^ and UDP formed well-diffracting crystals in the presence of the precipitant 2-methyl-2,4-pentanediol, with protein homodimers packing in a previously unreported C222_1_ lattice (**Figure 1f**; **Supplemental Figure 2**; **Supplemental Table 1**). Notably, the active sites of both PmHS2 catalytic domains appeared to be highly accessible in this new crystal form, with each domain containing bound UDP and Mn^2+^ cofactors, coordinating to characteristic GT DXD motifs^7^. Mn^2+^ ions in GlcA-T active sites were 5-coordinate with a distorted square pyramidal geometry, whilst Mn^2+^ in GlcNAc-T sites adopted 6-fold octahedral coordination (**Figure 1g, h**; **Supplemental Figure 3**).

### Structural mapping of the bifunctional PmHS2 reaction cycle

Having obtained a new, apparently ligandable form of PmHS2, we next explored structural characterization of its bifunctional reaction cycle, using alternating GlcNAc or GlcA terminated oligosaccharides to map heparosan binding across both enzyme active sites. Because PmHS2 catalyzes iterative elongation, we reasoned that the product of one catalytic domain would necessarily be the substrate of the other, enabling the start and end points of both transferase reactions to be mapped using only two ligands. Crystals of Δ80-PmHS2 were soaked with chemoenzymatically synthesized GlcNAc-GlcA-GlcNAc-GlcA-pNP (hereafter **4-mer**), or GlcA-GlcNAc-GlcA-GlcNAc-GlcA-pNP (**5-mer**)^28^. This approach produced well-resolved complexes containing unambiguous oligosaccharide densities across both GlcA-T and GlcNAc-T active sites of the enzyme, providing new insight into the full bifunctional PmHS2 reaction cycle (**Figure 2a**; **Supplemental Table 1**; **Supplemental Figure 4**).

**Figure 2.**
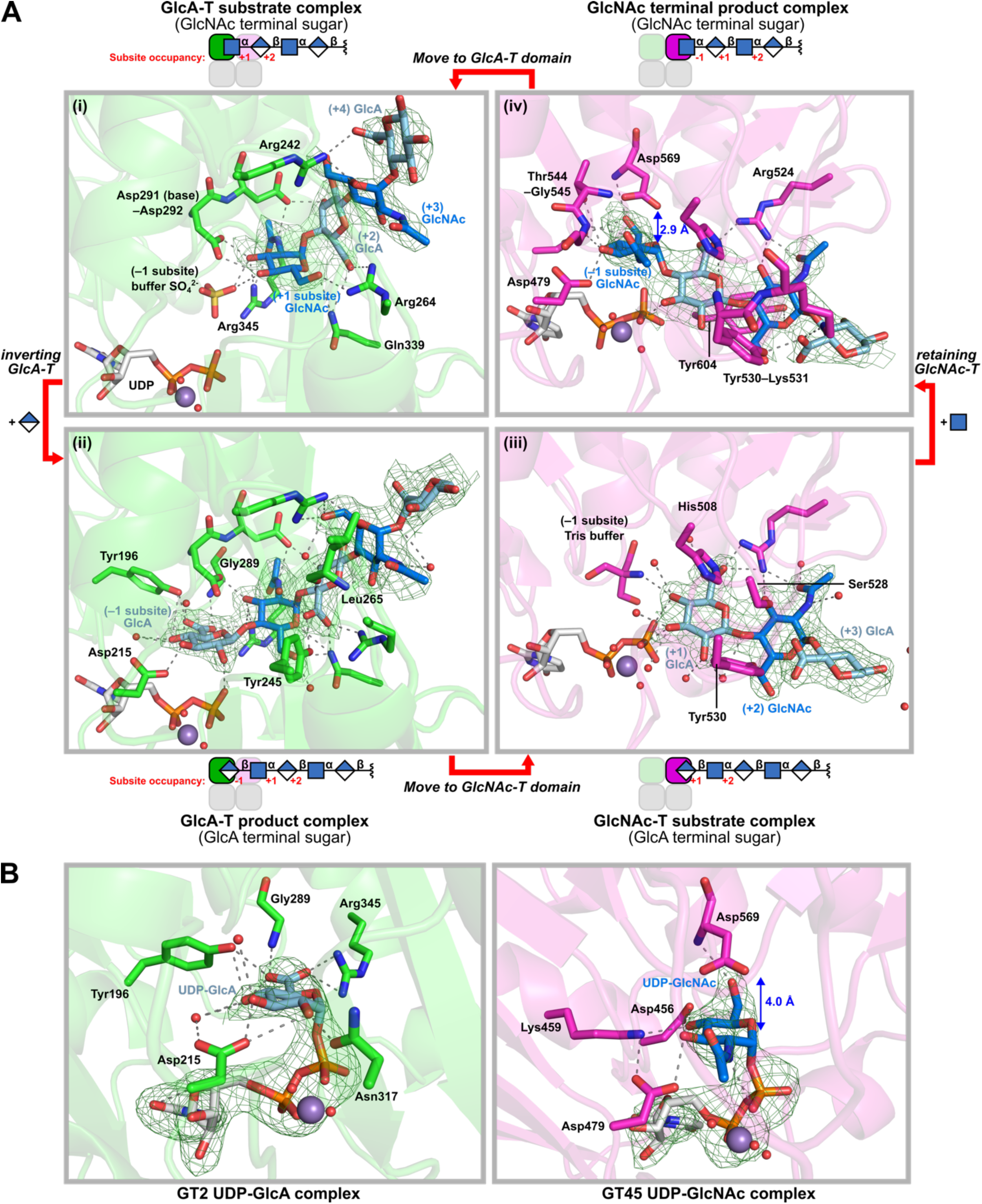
Acceptor and donor complexes map PmHS2 substrate-to-product trajectories, and suggest a covalent reaction pathway. (**A**) (i) **4-mer** and (ii) **5-mer** within the GlcA-T active site (green) delineate a reaction from substate to product. Conversely, **5-mer** (iii) and **4-mer** (iv) represent substrate and product for GlcNAc-T (pink). (**B**) Bound UDP-sugar donor substrates within the GlcA-T and GlcNAc-T active sites. For the GlcNAc-T domain, both donor substrate and acceptor product complexes show Asp5CS ‘in line’ with the anomeric center of GlcNAc, indicating a potential nucleophilic role. Electron densities are REFMAC σ_A_-weighted mFo–DFc, contoured between 2.0–2.4 σ (0.11–0.17 e^−^.Å^−3^).

First, within the inverting GlcA-T domain, inverting transfer was precisely described by its complex to substrate ligand **4-mer**, which occupied enzyme subsites +1 to +4 (numbering as in Ref. 29), and its complex to product ligand **5-mer**, which occupied –1 to +4 (**Figure 2a***i-ii*). These oligosaccharides bound the GlcA-T active site with offset register, such that the terminal non-reducing end (NRE) GlcNAc of **4-mer** was positioned at +1, and the NRE GlcA of **5-mer** at –1, delineating the start and end points of GlcA transfer onto a defined heparosan acceptor (**Supplemental Figure 5**). In both complexes, bound oligosaccharides were stabilized throughout the active site tunnel by multiple H-bonds, as well as by salt bridges from Arg242, Arg264 and Arg345 to the carboxylates of each GlcA. Close coordination between Asp291 to the 4OH nucleophile of NRE GlcNAc was observed in the **4-mer** substrate complex, confirming the essential role of this base within the inverting GT mechanism, which deprotonates GlcNAc to drive attack upon a bound UDP-GlcA donor (**Figure 1a**).

Next, retaining GlcNAc transfer within the PmHS2 GT45 domain was mapped in reciprocal fashion by its complexes to **5-mer** substrate and **4-mer** product, which bound with their NRE GlcA and GlcNAc residues at the enzyme +1 and –1 subsites, respectively (**Figure 2a***iii-iv*; **Supplemental Figure 5**). Strikingly, within the **4-mer** product complex, we observed the sidechain of Asp569 to sit directly ‘above’ the anomeric center of terminal NRE GlcNAc, poised for ‘backside’ attack, and suggesting possible mechanistic involvement during transfer. Although electrostatic stabilization of an oxocarbenium-like transition state can, in principle, be invoked for anionic sidechains within GT active sites, both the close distance (∼2.9 Å Asp569_OD2_-GlcNAc_C1_) and near in-line angle (158° ∠Asp569_OD2_-GlcNAc_C1_-GlcNAc_O1_) of Asp569 to GlcNAc were reminiscent of concerted nucleophilic attack, such as that required for covalent GT reactions.

Finally, to further complement structural analyses of PmHS2 acceptor complexes, we generated a co-complex of PmHS2 bound to both GlcA-T and GlcNAc-T glycosyl donors, by soaking crystals with an excess of both UDP-GlcNAc and UDP-GlcA ligands (**Figure 2b**). Within the resulting co-complex, each UDP-sugar was found to occupy its respective catalytic domain (UDP-GlcA in GlcA-T, UDP-GlcNAc in GlcNAc-T), *via* essentially the same Mn^2+^ coordination modes as previously observed for UDP. Notably, the glycosyl moieties of both UDP-GlcA and UDP-GlcNAc donors overlaid well with their corresponding heparosan product complexes, consistent with glycosyl transfer from these Michaelis complex positions onto approaching acceptor oligosaccharides (**Supplemental Figure 6**). As with the GlcNAc-T product complex (**Figure 2a***vi*), we again observed a close (∼4.0 Å), in-line placement between Asp569 and the anomeric center of UDP-GlcNAc donor, reinforcing a possible nucleophilic role for this sidechain during catalytic GlcNAc transfer.

### Computational evidence for a GT45 covalent intermediate

Observations of a poised Asp569 within the PmHS2 GlcNAc-T domain prompted us to further examine the role of this residue in catalytic transfer. Notably, amongst retaining hexosyltransferases, only enzymes from CAZy family GT6 have been previously proposed to harbor nucleophilic (glutamate) sidechains within their active sites^16^. Whilst investigations by us and others have suggested that GT6 enzymes can catalyze glycosyl transfer *via* covalent intermediate formation at these glutamates^14^, more passive roles for these residues in organizing pre-catalytic Michaelis complexes have also been proposed^17^.

When compared to GT6 enzymes (*e.g.* PDB 5NRD), Asp569 in PmHS2 appears even better placed for attack upon glycosyl donors, with its carboxylate located both closer to the UDP-sugar anomeric center (4.0 *vs* 4.4 Å in 5NRD), and along a more linear angle (158° *vs* 148°; **Supplemental Figure 7**). The close (2.9 Å) proximity of Asp569 to the GlcNAc-T product NRE is also unprecedented, and points to a tight pre-release complex following a second S_N_2 attack during transfer (**Figure 2a***vi*). To examine these PmHS2 mechanisms in detail, we employed quantitative Molecular Dynamics (MD) and Quantum Mechanics/Molecular Mechanics (QM/MM) methods, to simulate PmHS2 retaining GlcNAc transfer at an atomistic level^30^.

As a starting model for simulation, the three terminal non-reducing sugars from the GlcNAc-T **5-mer** substrate complex (**Figure 2a***iii*) were overlaid onto its UDP-sugar structure (**Figure 2b**), creating a fully occupied *in silico* substrate complex (**S**) containing both glycosyl donor and acceptor in the same active site (**Figure 3**). Classical MD simulations (three replicas 300 ns each) confirmed the stability of **S**, with C1_GlcNAc_–O4_acceptor_ and C1_GlcNAc_–Asp569 distances centering around ∼5.0 Å and ∼4.1 Å respectively (**Figure 3a**; **Supplemental Movie 1**). To investigate the endpoint of transfer, MD simulations were also conducted on the experimentally derived GlcNAc-T product complex with **4-mer** (**P**). Simulations again confirmed a stable complex, with C1_GlcNAc_–Asp569 distances centering around ∼3.3 Å (**Figure 3b**).

**Figure 3.**
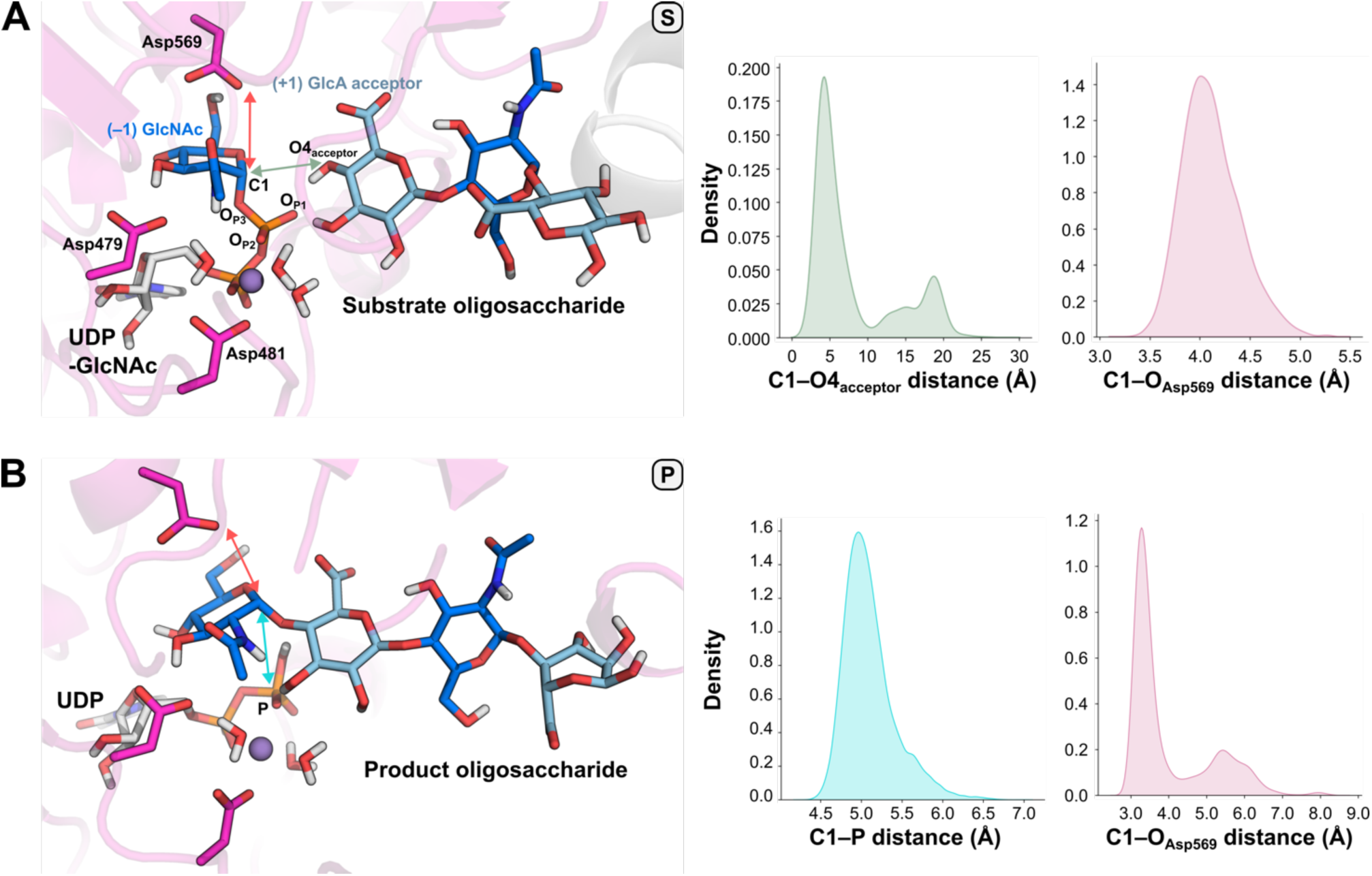
Classical MD simulations of GlcNAc-T starting and product complexes. (**A**) MD snapshot and distance histograms for the ternary substrate complex **S**, showing broad stability of the composite structure over 300 ns simulations. (**B**) MD snapshot and distance histograms for the product complex **P** also suggest stability.

With seemingly feasible **S** and **P** complexes confirmed, we progressed to QM/MM metadynamics simulations. As previously described for retaining GTs^14,31,32^, free energy landscapes of retaining GlcNAc transfer were calculated with respect to two collective variables (CVs), each summarizing defined mechanistic features of the system (*e.g.* key interatomic distances).

We first examined the possibility of (canonical) S_N_i-like transfer from **S**, using CVs optimized for this process, representing the initial substrate bond between GlcNAc and UDP β-phosphate (C1_GlcNAc_–O_P3_), and the final product bond between GlcNAc and GlcA (C1_GlcNAc_–O4_acceptor_). Notably, a distance between GlcNAc and Asp569 was not specified, to prevent bias towards a nucleophilic mechanism. Strikingly, this simulation did not evolve *via* an anticipated S_N_i-like trajectory, but instead proceeded to form a covalent glycosyl-enzyme intermediate (**GEI**) between GlcNAc and the sidechain of Asp569, suggesting that covalent, rather than S_N_i-like transfer may operate within the GlcNAc-T active site pocket (**Supplemental Figure 8**). To further test these striking observations, we reinitiated QM/MM using CVs explicit for covalent-intermediate formation, including the distance between GlcNAc and Asp569 (C1_GlcNAc_–O_D569_), alongside the C1_UDP-GlcNAc_–O_P3_ distance specified previously. Again, progression towards a GlcNAc-linked **GEI** was observed, *via* a concerted mechanism with a computed free energy barrier of 19.0 kcal/mol (**Supplemental Figure 9**), in good agreement with values estimated from prior kinetics experiments (k_cat_ = 3.5 s^-1^; 17.1 kcal/mol)^22^.

Despite extensive optimization, we were unable to simulate full transfer from **S** to form glycosyltransferase reaction products, instead consistently observing re-equilibration between **GEI** and **S**. We considered this may be due to the origin of **S** as an *in silico* composite of UDP-sugar and heparosan structures, which does not reflect binding within a true Michaelis complex. QM/MM simulations were thus reinitiated from product complex **P**, using this experimentally derived model to calculate a reverse transfer reaction. Intriguingly, **P** also reacted *in silico* to form a **GEI** containing Asp569 bound GlcNAc (**Figure 4a**), and, moreover, could be regressed to a substrate state (**S’** to distinguish from composite complex **S**) using a second set of CVs representing C1_GlcNAc_–P and C1_GlcNAc_– O_D569_ (**Figure 4b**). **S’** differs from **S** by a closer positioning of heparosan acceptor substrate to the UDP-GlcNac donor (**Supplemental Figure 10**), suggesting that initial ternary complexes (*e.g.* **S**) may evolve towards more catalytically competent states, in a fashion not captured by MD. Because the principle of microscopic reversibility applies to enzymatic reactions^33^, we consider simulations from **P** to **S’** to be valid descriptors of GlcNAc transfer across both forwards and reverse directions, and thus **S’** to be a predictive Michaelis complex for the forward transfer reaction.

**Figure 4.**
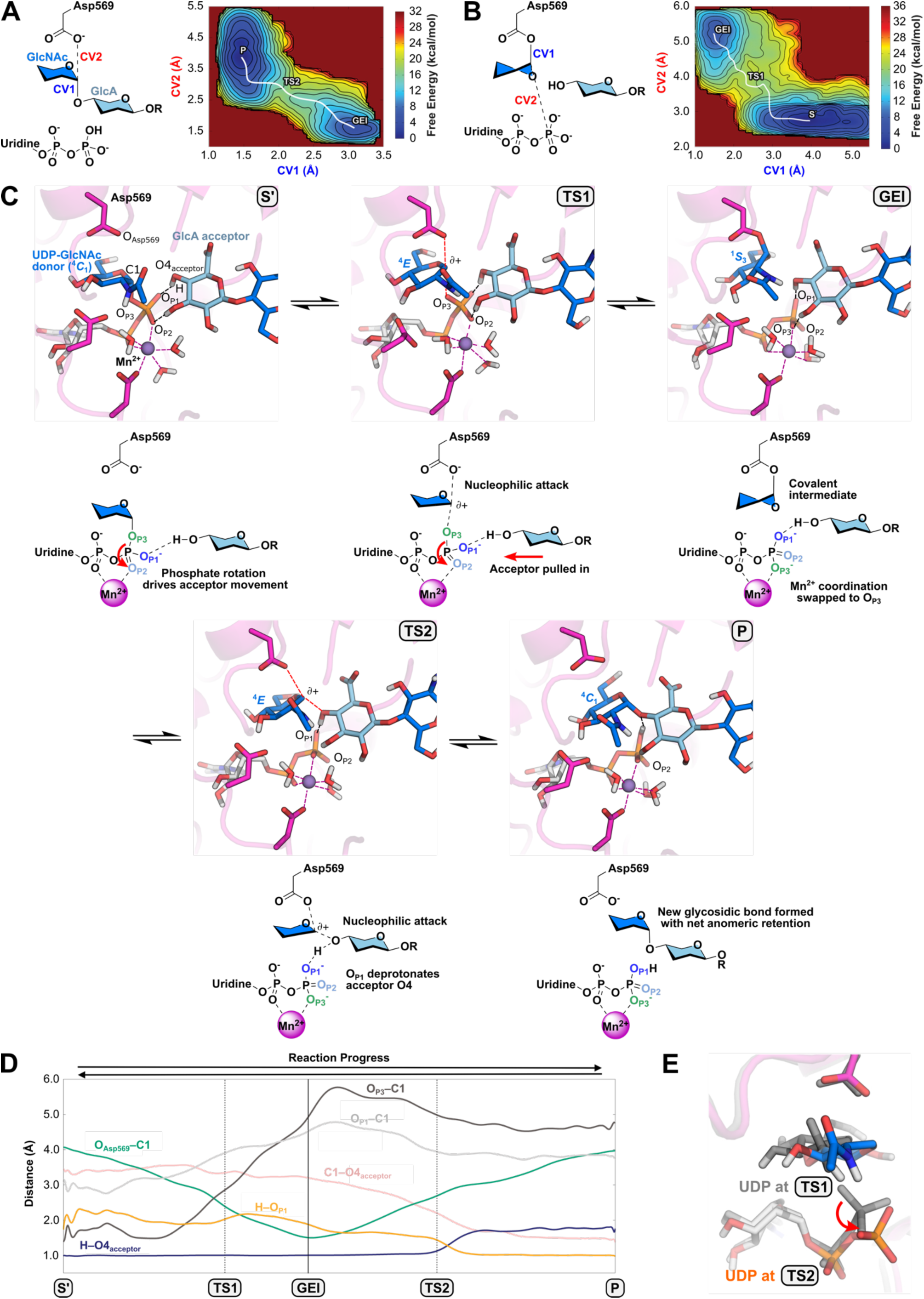
QM/MM simulations of the retaining GlcNAc-T reaction. CVs and FELs for reverse reaction from **P** to **GEI** (**A**) and **GEI** to **S’** (**B**). (**C**) Snapshots of the transfer from **S’** to **P**, via formation of a covalent **GEI**. Initial nucleophilic attack upon the UDP-GlcNAc anomeric center is accompanied by rotation of the UDP pyrophosphate, which pulls the acceptor sugar towards the reacting donor. Deprotonation of the acceptor by UDP within the active site drives attack upon **GEI**, leading to its breakdown and formation of the GlcNAc-T product. (**D**) Changes to selected bond lengths from **S’** to **P**. (**E**) UDP β-phosphate positions at **TS1** and **TS2**, highlighting rotational movement.

Detailed examination of simulated reaction trajectories showed that **S’** to **GEI** proceeds *via* initial extension of the UDP-GlcNAc sugar-phosphate bond (C1_GlcNAc_–O_P3_), driving conformational distortion of GlcNAc from a ^4^*C*_1_ chair to a ^4^*E* envelope at its oxocarbenium-like transition state (**TS1**), crossing a free energy barrier of 21.2 kcal/mol (**Figure 4c, d**). From **TS1**, where GlcNAc is essentially equidistant between Asp569 and UDP (2.7 vs 2.8 Å), further movement towards Asp569 results in formation of the **GEI**, with the enzyme-linked sugar adopting a ^1^*S*_3_ skew-boat conformation, reminiscent of retaining glycoside hydrolases^34^. At the **GEI**, O4 of the acceptor GlcA is also positioned ∼5 Å from the bound GlcNAc intermediate C1, with a calculated nucleophilic approach angle of ∼130°.

Finally, transition from **GEI** towards **P** proceeds with elongation of the C1_GlcNAc_–O_D569_ bond, allowing detachment of GlcNAc to initiate its movement towards the acceptor NRE GlcA. This second transfer also proceeds through an oxocarbenium-like transition state (**TS2**; 12.7 kcal/mol), with GlcNAc again adopting a distorted ^4^*E* conformation. Formation of the product C1_GlcNAc_–O4_acceptor_ bond concludes with attack of the acceptor O4 onto **TS2**, supported by concurrent abstraction of the OH proton by a basic phosphate from departing UDP (**Figure 4c**,**d**). The full reaction from **S’** to **P** is depicted in **Supplemental Movie 2**.

### Phosphate rotation as a geared base mechanism for transfer

A further intriguing aspect of PmHS2 retaining GlcNAc transfer revealed by our QM/MM simulations is a large movement of the UDP β-phosphate, which rotates by ∼60° as the reaction progresses from **S’** to **GEI** – a mechanistic feature not seen in other retaining GTs (**Figure 4e**). Due to this rotation, O_P3_ (the phosphate oxygen initially linked to GlcNAc) moves closer to Mn^2+^ as the reaction proceeds, eventually displacing O_P2_ to coordinate the Mn^2+^ center (**Figure 4c,d**). This rotational motion also causes O_P1_ to pull on the acceptor NRE sugar, enabling its O4_acceptor_ moiety to approach the covalent **GEI**, before subsequent deprotonation to initiate nucleophilic attack. The GlcNAc-T reverse reaction is accompanied by retrograde β-phosphate rotation, which pushes the acceptor heparosan away as UDP-GlcNAc reforms. We propose that phosphate rotation, coupled to Mn^2+^ coordination, may act as a ‘geared base’ in PmHS2 GlcNAc-T reactions, organizing transfer across both forwards or backwards directions, depending on its overall direction of movement.

### Intrinsic nucleophilicity of GlcNAc-T active site position 5CS

Given the strong computational evidence for a covalent intermediate during PmHS2 GlcNAc transfer, we next sought to verify this mechanism by capturing its glycosyl-enzyme intermediate. As wild-type Δ80-PmHS2 may react too fast to detect any intermediate bound to Asp569, we generated putative nucleophile mutants D569N and D569C to trap potential adducts, as well as D569A, which should lack reactivity. Strikingly, intact mass-spectrometry of Δ80-PmHS2-D569C directly following purification revealed an unexpected +203 Da mass shift, consistent with addition of a GlcNAc following reaction with intracellular UDP-GlcNAc (**Figure 5a**). By contrast, treatment of purified wild-type, D569A, or D569N Δ80-PmHS2 with 10 mM UDP-sugars for 10 minutes did not produce any observable mass shifts (**Supplemental Figure 11**).

**Figure 5.**
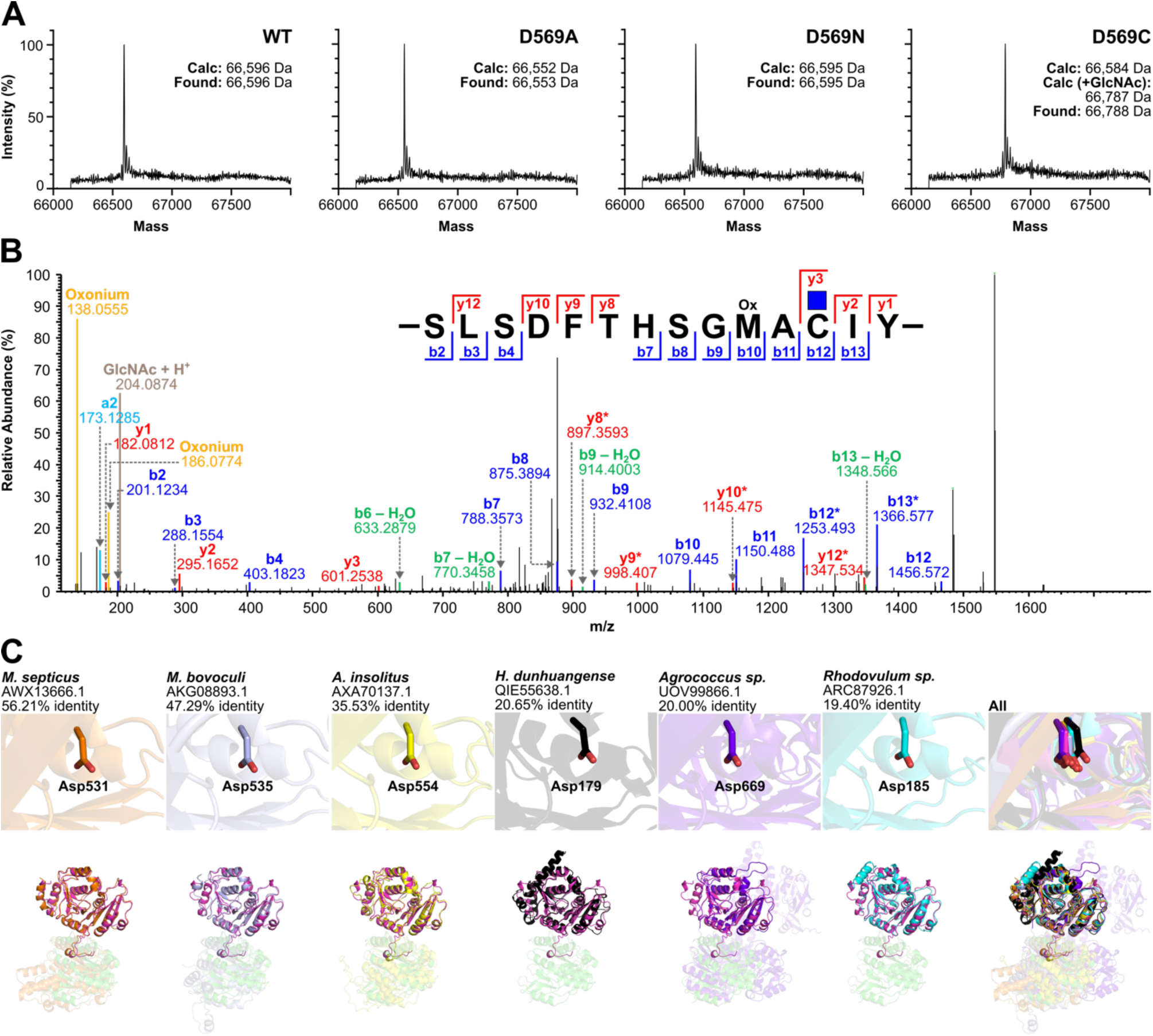
Nucleophilicity of position 5CS in PmHS2 and GT45 homologs. (**A**) ESI-MS spectra of purified Δ80-PmHS2 wild-type and D5CSA, D5CSN and D5CSC mutants. A +203 Da mass shift was observed for D5CSC, consistent with addition of GlcNAc. (**B**) MS/MS sequencing of a chymotryptic peptide confirms the presence of a HexNAc adduct on the mutated Cys5CS residue. (**C**) AF3 models of PmHS2 homologs, showing near universal conservation of the Asp5CS equivalent position across the CAZy GT45 family. Top – active sites. Bottom – 3-dimensional folds of GT45 domains, overlaid onto PmHS2 (pink).

To confirm if the mass shift on Δ80-PmHS2-D569C resulted from GlcNAc addition to its mutated cysteine, we first attempted X-ray crystallography. Additional electron density could be observed around the Cys569 position of Δ80-PmHS2-D569C in its crystal structure, but the marginal ∼3.9 Å resolution of this dataset prevented confident assignment (**Supplemental Figure 12**). Mass-spectrometry (MS)-based sequencing was thus employed for further verification. Analysis of a chymotryptic peptide around the D569C mutation site revealed the clear presence of a +203 Da adduct, which was confirmed by MS2 to reside on Cys569. Notably, whilst Cys mutants are often used to probe nucleophilicity of active site positions in GTs and other enzymes, trapping experiments typically require substrates to be added *in vitro* before adducts can be observed^12,13^. Our observation that PmHS2-D569C forms GlcNAc adducts directly from intracellular milieu suggests a highly favorable interaction, underscoring likely intrinsic nucleophilicity of this position within the overall enzyme fold (**Figure 5b**).

Finally, we considered whether our observed nucleophilicity may be a general characteristic of the broader GT45 family to which the PmHS2 GlcNAc-T domain belongs. Clustal Omega^35^ alignment of 51 representative GT45 sequences from CAZy (1 sequence per genus) indicated that the Asp569 position is almost fully conserved, with the sole exception being a cryptic GT45 glycosyltransferase from Sulfitobacter, which also lacks a DXD motif (**Supplemental Figure 13**). Comparison of PmHS2 to AlphaFold3^36^ predictions of its 3 most and 3 least homologous sequences showed full structural conservation of the Asp569 equivalent position (**Figure 5c**), supporting an essential role for this residue during catalysis, and likely conservation of covalent mechanisms across the GT45 family.

### PmHS2 self-priming via interplay of UDP-sugar interactions

Bifunctional glycosyltransferases such as PmHS2 must coordinate interactions between two sets of substrates. Whilst such interactions are typically independent, an unusual example of functional cooperativity is seen in the self-priming reaction of PmHS2, whereby the enzyme initiates polymerization of new oligosaccharides upon incubation with UDP-GlcA and UDP-GlcNAc donors^37^ (**Figure 1c, d**). In our crystallographic study, we noted clear additional density within the GT45 active site following short (<5 min) UDP-sugar co-incubations, which correlated well with **5-mer** complexes in this same domain (**Supplemental Figure 14**).

Inspection of the PmHS2 UDP-sugar ternary complex revealed a likely mechanism for *de novo* polymerization. Unexpectedly, the PmHS2 GlcA-T domain was found to carry a previously unreported secondary UDP-GlcNAc binding site, centered around a surface Mn^2+^ ion coordinated to both Asp292 and His296 (**Figure 6a, b**). UDP-GlcNAc was found to bind this secondary site such that its sugar moiety occupied the GlcA-T +1 region, coordinated to Asp291 in a similar fashion to the **4-mer** NRE – poised for attack upon a bound UDP-GlcA donor (**Figure 6c**). Consequently, UDP-GlcNAc appears capable of functioning as a ‘mini-primer’ within the GlcA-T domain, which can substitute for *bona fide* heparosan acceptors in cases of their absence; no similar secondary UDP-sugar site could be identified in the GlcNAc-T domain. Taken together, our observations strongly indicate that PmHS2 *de novo* polymerization occurs at its GT2 domain, *via* an initial attack of UDP-GlcNAc onto a UDP-GlcA donor. These structures are in line with prior biochemical studies of PmHS2 self-priming, including reports that newly initiated polysaccharides contain attached UDP moieties^38^ (**Figure 6d**). PmHS2 non-canonical substrate binding likely provides an effective mechanism to initiate heparosan polymerization, and represents the first (to our knowledge) example of a GT using intrinsically electrophilic glycosyl donors also as nucleophilic glycosyl acceptors.

**Figure 6.**
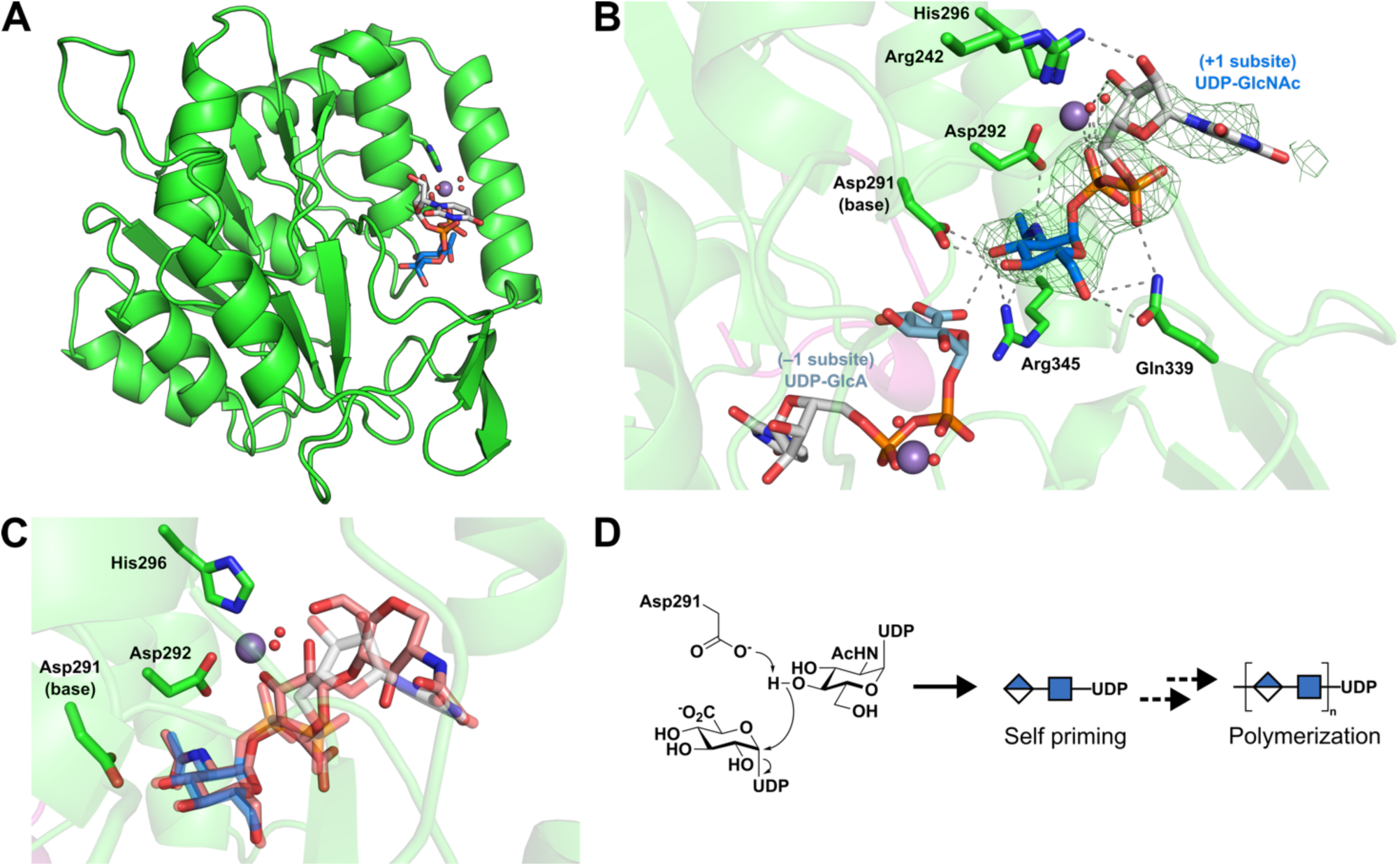
Structural basis of PmHS2 self-priming. (**A**) Surface localized binding of a secondary UDP-GlcNAc at the GlcA-T domain. (**B**) Close-up view of secondary UDP-GlcNAc binding with respect to UDP-GlcA donor. GlcNAc interacts with the Asp291 base as would be expected of a glycosyl acceptor nucleophile. (**C**) Overlay of secondary UDP-GlcNAc binding compared to **4-mer** substrate binding, showing its pseudo-mimicry of an acceptor oligosaccharide. (**D**) Mechanistic proposal for GlcA-T self-priming, via initial attack of a UDP-GlcNAc acceptor onto a UDP-GlcA donor.

## Discussion

Precise analysis of GT mechanisms is essential to understanding biological control of glycosylation, and paves the way for their modulation in biomedical or biotechnological applications. Here, we conducted a comprehensive structure/function study of bifunctional PmHS2, which contains both inverting GlcA-T and retaining GlcNAc-T activities within the same protein chain. Mapping substrate and product binding modes across both PmHS2 catalytic domains provided key insight into their mechanistically distinct active sites, building considerably upon previous reports of the enzyme 3-dimensional fold^18^.

Mechanistically, the GlcA-T domain of PmHS2, belonging to CAZy family GT2, catalyzes inverting transfer of GlcA from UDP-linked donors onto heparosan acceptors bearing NRE GlcNAc sugars. Our experimentally obtained complexes with substrate **4-mer** and product **5-mer** are consistent with the known activity of this domain. Specifically, we find that the catalytic base Asp291 closely coordinates with O4 of the acceptor GlcNAc within its substrate complex, enabling facile deprotonation to assist attack upon an active-site UDP-GlcA.

Alongside the PmHS2 GlcA-T domain, a separate retaining GlcNAc-T domain (CAZy family GT45) catalyzes transfer of GlcNAc onto heparosan acceptors bearing NRE GlcA sugars. Our combined structural and computational study here represents the first detailed investigation of this retaining GT family, revealing a transfer mechanism involving likely covalent adduct formation to the sidechain of Asp569. PmHS2 thus joins the retaining Kdo-transferases WbbB and KpsC^12,13,39,40^, and the transferases from family GT6^14^, as one of the few retaining GTs proposed to employ a covalent reaction intermediate.

Despite superficial similarities regarding their covalent transfer reactions, important mechanistic distinctions may be made between the retaining Kdo octulosyltransferases, and the hexosyltransferases from GT6 and GT45 (*i.e.* PmHS2). Notably, retaining Kdo-transferases are reported to employ a ‘ping-pong’-like reaction pathway involving initial formation of a glycosyl-enzyme intermediate with loss of nucleotide aglycone, followed by deprotonation of an incoming glycosyl acceptor by an enzyme-contributed base. This process bears close similarity to retaining glycoside hydrolase mechanisms, wherein the enzyme active site processes two distinct half reactions, in which departing aglycone is replaced by an incoming nucleophilic water. By contrast, both GT6 and GT45 hexosyltransferases appear to proceed *via* a single pre-organized active site, wherein the departing sugar nucleotide aglycone (UDP) remains associated with the reaction coordinate throughout. These reactions may, in principle, be considered as an extreme example of ion-pair stabilization between the positive oxocarbenium-like glycosyl intermediate and the negatively charged enzyme carboxylate, wherein sufficiently close contact is achieved to enable covalent bond formation. Our results, alongside prior observations from GT6, therefore suggest that covalent GT mechanisms may lie along a continuum, spanning from reactions more similar to classical S*_N_*i transfer (such as here), to those that employ hydrolase-like double displacement (*i.e.* Kdo-transferases).

Beyond formation of a covalent adduct, a further notable feature observed during PmHS2 GlcNAc transfer was its UDP β-phosphate rotation, which appears to direct reaction progress *via* ‘geared’ coordination of the active site Mn^2+^. Notably, compared to bidentate protein sidechains that typically coordinate to metals (e.g. Asp/Glu), the tridentate geometry of UDP phosphates may offer GTs the capability to interact with metal, sugar, and acceptor nucleophiles, all from a single center, providing additional utility for multicomponent reactions. By coupling UDP phosphate rotation to oligosaccharide movement, the PmHS2 acceptor substrate may be readily moved in or out of the enzyme catalytic center, driving reactivity against trapped glycosyl-enzyme intermediates. Our observations hint at hitherto underappreciated abilities of nucleotide cofactors to direct GT reactions as geared ‘mechanical devices’, in addition to their more common ‘electronic’ roles as activated leaving groups and bases.

Finally, by generating PmHS2 ligand complexes with UDP-sugar acceptors, we also identified an unexpected mechanistic basis for its self-priming activity, which has (in part) led to the adoption of this enzyme for chemoenzymatic heparosan production^20,21^. Mn^2+^ driven binding of UDP-GlcNAc to a secondary site on the PmHS2 GlcA-T domain allows for this sugar nucleotide to act as a glycosyl acceptor (rather than donor) at this site, effectively substituting for longer heparosan primers to initiate attack upon UDP-GlcA. Identifying structural principles that underpin *de novo* polymerization may enable similar functions to be engineered into polymerases that lack self-priming, such as EXT1/2^24^, PmHS1^41^, or PmHAS^42^, broadening their utility for chemoenzymatic exploitation.

In summary, our results highlight the substantial mechanistic insights that are still to be discovered within the GT class of enzymes. Understanding GT enzyme-substrate binding, transition states, and conformational itineraries will underpin the rational development of inhibitors^43,44^, or the engineering of novel activities^45,46^. The finding of a key covalent intermediate in PmHS2 GlcNAc transfer highlights a pathway that is exploitable through chemical design, opening the door to bespoke routes for modulating this and related GTs.

## Data availability

All structures generated in this study have been deposited in the PDB under accession codes 34ZL (PmHS2-Mn^2+^-UDP complex), 34ZN (PmHS2-Mn^2+^-UDP-4-mer complex), 34ZM (PmHS2-Mn^2+^-UDP-5-mer complex), 34ZO (PmHS2-Mn^2+^-UDP-sugar co-complex) and 34ZP (PmHS2-D569C). MS/MS sequencing data for the PmHS2-D569C-GlcNAc adduct have been deposited in PRIDE, under accession code PXD078915.

## Author contributions

MS, BP, CR – simulations

MAL, ESK, SM – intact and sequencing MS

CJMW – enzyme kinetics

GP, MAG, LW – cloning and purification

LW – crystallography

BGD, CR, LW – conception, supervision, writing

## Acknowledgements

The Rosalind Franklin Institute is funded by UK Research and Innovation through the Engineering and Physical Sciences Research Council (UKRI-EPSRC). The Chemistry theme at the Rosalind Franklin Institute has been further supported by grants from UKRI-EPSRC (EP/V011359/1, EP/V011356/1, EP/T012021/1, EP/X527245/1). GP was supported by an Onassis Foundation PhD scholarship (F ZQ 048-1/2020-2021) and a St Edmund Hall scholarship (William R Miller Postgraduate Award 2020-2021). LW and CMW acknowledge support by the Wellcome Trust (Sir Henry Dale Fellowship 218579/Z/19/Z to LW). CR acknowledges funding from the Spanish Ministry of Science, Innovation and Universities (MICIU/AEI/10.13039/501100011033, ref. PID2023-147939NB-100) and the Spanish Structures of Excellence María de Maeztu (CEX2021-001202-M). We are grateful for the technical assistance provided by the support of the MareNostrum IV of the Barcelona Supercomputing Center (BSC-CNS). We thank Diamond Light Source for access to beamlines I03 and I04 (proposals mx-34263 and mx-28534), which contributed to the results presented here.

## Supporting information

The authors declare no competing financial interests.

## References

1. . Egan, A. J. F.; Errington, J.; Vollmer, W., Nat Rev Microbiol 2020, 18 (8), 446–460.

2. . Scheller, H. V.; Ulvskov, P., Annu Rev Plant Biol 2010, 61, 263–89.

3. . Pellegrini, L., Curr Opin Struct Biol 2001, 11 (5), 629–34.

4. . Crocker, P. R.; Paulson, J. C.; Varki, A., Nat Rev Immunol 2007, 7 (4), 255–66.

5. . Dedola, S.; Rugen, M. D.; Young, R. J.; Field, R. A., ChemBioChem 2020, 21 (3), 423–427.

6. . Lombard, V.; Golaconda Ramulu, H.; Drula, E.; Coutinho, P. M.; Henrissat, B., Nucleic Acids Res 2014, 42 (Database issue), D490–5.

7. . Lairson, L. L.; Henrissat, B.; Davies, G. J.; Withers, S. G., Annu Rev Biochem 2008, 77, 521–55.

8. . Ardevol, A.; Rovira, C., J Am Chem Soc 2015, 137 (24), 7528–47.

9. . Lee, S. S.; Hong, S. Y.; Errey, J. C.; Izumi, A.; Davies, G. J.; Davis, B. G., Nat Chem Biol 2011, 7 (9), 631–8.

10. . Ardevol, A.; Rovira, C., Angew Chem Int Ed Engl 2011, 50 (46), 10897–901.

11. . Davies, G.; Henrissat, B., Structure 1995, 3 (9), 853–9.

12. . Doyle, L.; Ovchinnikova, O. G.; Huang, B. S.; Forrester, T. J. B.; Lowary, T. L.; Kimber, M. S.; Whitfield, C., J Biol Chem 2023, 299 (5), 104609.

13. . Forrester, T. J. B.; Ovchinnikova, O. G.; Li, Z.; Kitova, E. N.; Nothof, J. T.; Koizumi, A.; Klassen, J. S.; Lowary, T. L.; Whitfield, C.; Kimber, M. S., Nat Commun 2022, 13 (1), 6277.

14. . Rojas-Cervellera, V.; Ardevol, A.; Boero, M.; Planas, A.; Rovira, C., Chemistry 2013, 19 (42), 14018–23.

15. . Blackler, R. J.; Gagnon, S. M.; Polakowski, R.; Rose, N. L.; Zheng, R. B.; Letts, J. A.; Johal, A. R.; Schuman, B.; Borisova, S. N.; Palcic, M. M.; Evans, S. V., Glycobiology 2017, 27 (4), 370–380.

16. . Gomez, H.; Lluch, J. M.; Masgrau, L., J Am Chem Soc 2013, 135 (18), 7053–63.

17. . Albesa-Jove, D.; Sainz-Polo, M. A.; Marina, A.; Guerin, M. E., Angew Chem Int Ed Engl 2017, 56 (47), 14853–14857.

18. . Stancanelli, E.; Krahn, J. A.; Viverette, E.; Dutcher, R.; Pagadala, V.; Borgnia, M. J.; Liu, J.; Pedersen, L. C., ACS Catalysis 2024, 6577–6588.

19. . Deangelis, P. L.; White, C. L., J Bacteriol 2004, 186 (24), 8529–32.

20. . Liu, J.; Linhardt, R. J., Nat Prod Rep 2014, 31 (12), 1676–85.

21. . Xu, Y.; Masuko, S.; Takieddin, M.; Xu, H.; Liu, R.; Jing, J.; Mousa, S. A.; Linhardt, R. J.; Liu, J., Science 2011, 334 (6055), 498–501.

22. . Na, L.; Yu, H.; McArthur, J. B.; Ghosh, T.; Asbell, T.; Chen, X., ACS Catal 2020, 10 (11), 6113–6118.

23. . Li, H.; Chapla, D.; Amos, R. A.; Ramiah, A.; Moremen, K. W.; Li, H., Nat Chem Biol 2023, 19 (5), 565–574.

24. . Leisico, F.; Omeiri, J.; Le Narvor, C.; Beaudouin, J.; Hons, M.; Fenel, D.; Schoehn, G.; Coute, Y.; Bonnaffe, D.; Sadir, R.; Lortat-Jacob, H.; Wild, R., Nat Commun 2022, 13 (1), 7110.

25. . Dutta, P.; Cordeiro, R. L.; Friedel-Arboleas, M.; Bourgeais, M.; Vallet, S. D.; Weber, M.; Molinas, M.; Shu, H.; Grønset, M. N. N.; Miller, R. L.; Boeri Erba, E.; Wild, R., Nature Communications 2025, 16 (1), 11663.

26. . Wilson, L. F. L.; Dendooven, T.; Hardwick, S. W.; Echevarría-Poza, A.; Tryfona, T.; Krogh, K. B. R. M.; Chirgadze, D. Y.; Luisi, B. F.; Logan, D. T.; Mani, K.; Dupree, P., Nature Communications 2022, 13 (1), 3314.

27. . Wang, Z.; Ly, M.; Zhang, F.; Zhong, W.; Suen, A.; Hickey, A. M.; Dordick, J. S.; Linhardt, R. J., Biotechnology and Bioengineering 2010, 107 (6), 964–973.

28. . DeAngelis, P. L.; Liu, J.; Linhardt, R. J., Glycobiology 2013, 23 (7), 764–777.

29. . Davies, G. J.; Wilson, K. S.; Henrissat, B., Biochem J 1997, 321 *( Pt* *2*) (Pt 2), 557–9.

30. . van der Kamp, M. W.; Mulholland, A. J., Biochemistry 2013, 52 (16), 2708–28.

31. . Lira-Navarrete, E.; Iglesias-Fernandez, J.; Zandberg, W. F.; Companon, I.; Kong, Y.; Corzana, F.; Pinto, B. M.; Clausen, H.; Peregrina, J. M.; Vocadlo, D. J.; Rovira, C.; Hurtado-Guerrero, R., Angew Chem Int Ed Engl 2014, 53 (31), 8206–10.

32. . Bilyard, M. K.; Bailey, H. J.; Raich, L.; Gafitescu, M. A.; Machida, T.; Iglesias-Fernandez, J.; Lee, S. S.; Spicer, C. D.; Rovira, C.; Yue, W. W.; Davis, B. G., Nature 2018, 563 (7730), 235–240.

33. . Astumian, R. D., Nat Nanotechnol 2012, 7 (11), 684–8.

34. . Speciale, G.; Thompson, A. J.; Davies, G. J.; Williams, S. J., Curr Opin Struct Biol 2014, 28, 1–13.

35. . Sievers, F.; Higgins, D. G., Curr Protoc Bioinformatics 2014, 48, 3 13 1-3 13 16.

36. . Abramson, J.; Adler, J.; Dunger, J.; Evans, R.; Green, T.; Pritzel, A.; Ronneberger, O.; Willmore, L.; Ballard, A. J.; Bambrick, J.; Bodenstein, S. W.; Evans, D. A.; Hung, C. C.; O’Neill, M.; Reiman, D.; Tunyasuvunakool, K.; Wu, Z.; Zemgulyte, A.; Arvaniti, E.; Beattie, C.; Bertolli, O.; Bridgland, A.; Cherepanov, A.; Congreve, M.; Cowen-Rivers, A. I.; Cowie, A.; Figurnov, M.; Fuchs, F. B.; Gladman, H.; Jain, R.; Khan, Y. A.; Low, C. M. R.; Perlin, K.; Potapenko, A.; Savy, P.; Singh, S.; Stecula, A.; Thillaisundaram, A.; Tong, C.; Yakneen, S.; Zhong, E. D.; Zielinski, M.; Zidek, A.; Bapst, V.; Kohli, P.; Jaderberg, M.; Hassabis, D.; Jumper, J. M., Nature 2024, 630 (8016), 493–500.

37. . Sismey-Ragatz, A. E.; Green, D. E.; Otto, N. J.; Rejzek, M.; Field, R. A.; DeAngelis, P. L., J Biol Chem 2007, 282 (39), 28321–28327.

38. . Chavaroche, A. A. E.; van den Broek, L. A. M.; Boeriu, C.; Eggink, G., Applied Microbiology and Biotechnology 2012, 95 (5), 1199–1210.

39. . Rao, D.; Zhu, L.; Liu, W.; Guo, Z., J Phys Chem B 2024.

40. . Sagiroglugil, M.; Liao, Q.; Planas, A.; Rovira, C., ChemCatChem 2024, n/a (n/a), e202400769.

41. . Otto, N. J.; Green, D. E.; Masuko, S.; Mayer, A.; Tanner, M. E.; Linhardt, R. J.; DeAngelis, P. L., J Biol Chem 2012, 287 (10), 7203–12.

42. . Williams, K. J.; Halkes, K. M.; Kamerling, J. P.; DeAngelis, P. L., J Biol Chem 2006, 281 (9), 5391–7.

43. . Artola, M.; Wu, L.; Ferraz, M. J.; Kuo, C. L.; Raich, L.; Breen, I. Z.; Offen, W. A.; Codee, J. D. C.; van der Marel, G. A.; Rovira, C.; Aerts, J.; Davies, G. J.; Overkleeft, H. S., ACS Cent Sci 2017, 3 (7), 784–793.

44. . de Boer, C.; Armstrong, Z.; Lit, V. A. J.; Barash, U.; Ruijgrok, G.; Boyango, I.; Weitzenberg, M. M.; Schroder, S. P.; Sarris, A. J. C.; Meeuwenoord, N. J.; Bule, P.; Kayal, Y.; Ilan, N.; Codee, J. D. C.; Vlodavsky, I.; Overkleeft, H. S.; Davies, G. J.; Wu, L., Proc Natl Acad Sci U S A 2022, 119 (31), e2203167119.

45. . Coines, J.; Cuxart, I.; Teze, D.; Rovira, C., J Phys Chem B 2022, 126 (4), 802–812.

46. . Iglesias-Fernandez, J.; Hancock, S. M.; Lee, S. S.; Khan, M.; Kirkpatrick, J.; Oldham, N. J.; McAuley, K.; Fordham-Skelton, A.; Rovira, C.; Davis, B. G., Nat Chem Biol 2017, 13 (8), 874–881.

