## Supplemental Information file for "Hexosyl transfer *via* a covalent intermediate within a dual retaining and inverting enzyme"

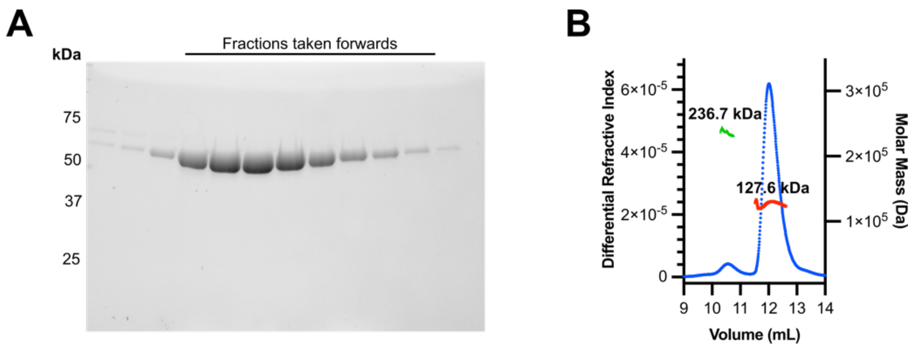


***Supplemental Figure 1*** *(**A**) SDS-PAGE of Δ80-PmHS2 after SEC purification. Protein chain shows monomeric mass of ~67 kDa. (**B**) SEC-MALS of purified Δ80-PmHS2 showing predominantly dimeric stoichiometry.*

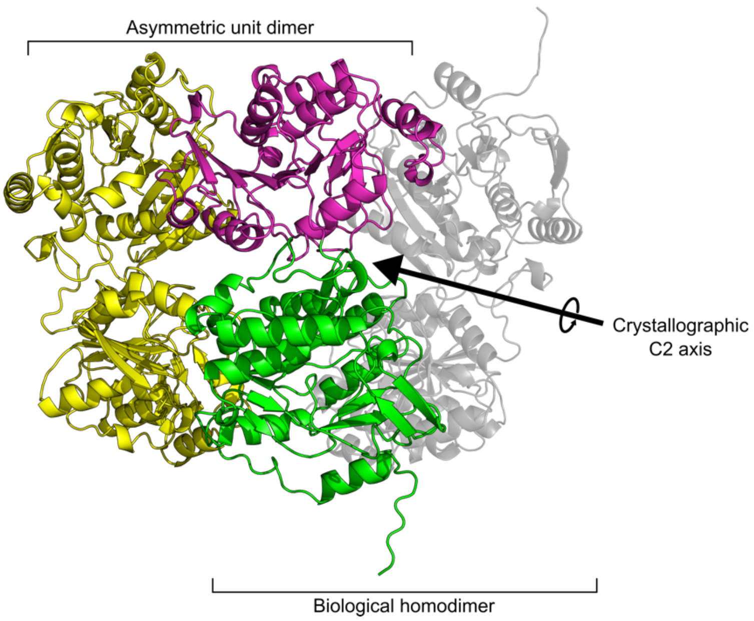


***Supplemental Figure 2*** *Molecular packing of the C222$_1$ crystal form of Δ80-PmHS2. Each asymmetric unit contains two molecules of Δ80-PmHS2 that form a non-biological dimer (monomer 1 - green/pink, monomer 2 – yellow). The biological homodimer (grey transparent) is formed by a C2 symmetry operation along a crystallographic axis.*

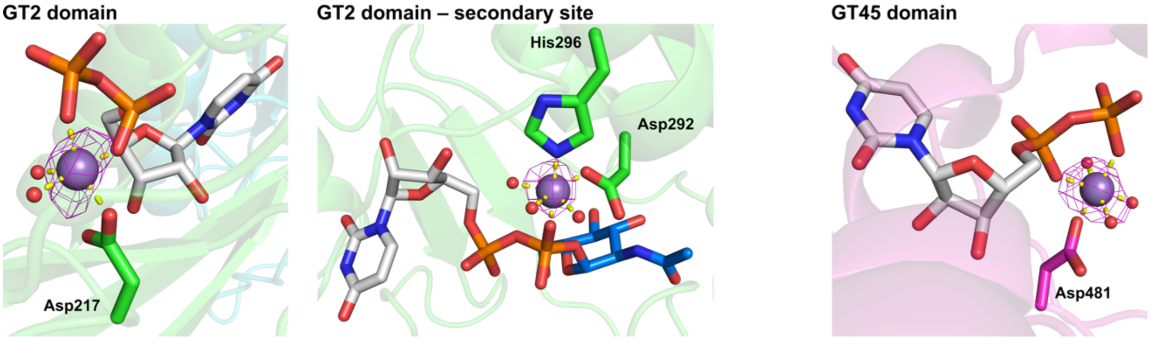


***Supplemental Figure 3*** *Comparison of $Mn^{2+}$ coordination across the PmHS2 structure. The primary GT2 and GT45 $Mn^{2+}$ sites are 5-coordinate and 6-coordinate, respectively. A secondary 6-coordinate $Mn^{2+}$ site in the GT2 domain aids UDP-GlcNAc binding to initiate de novo polymerization. Densities are anomalous maps to confirm the presence of $Mn^{2+}$, contoured to 3–5 σ (0.03–0.05 $e^{-}.Å^{-3}$).*

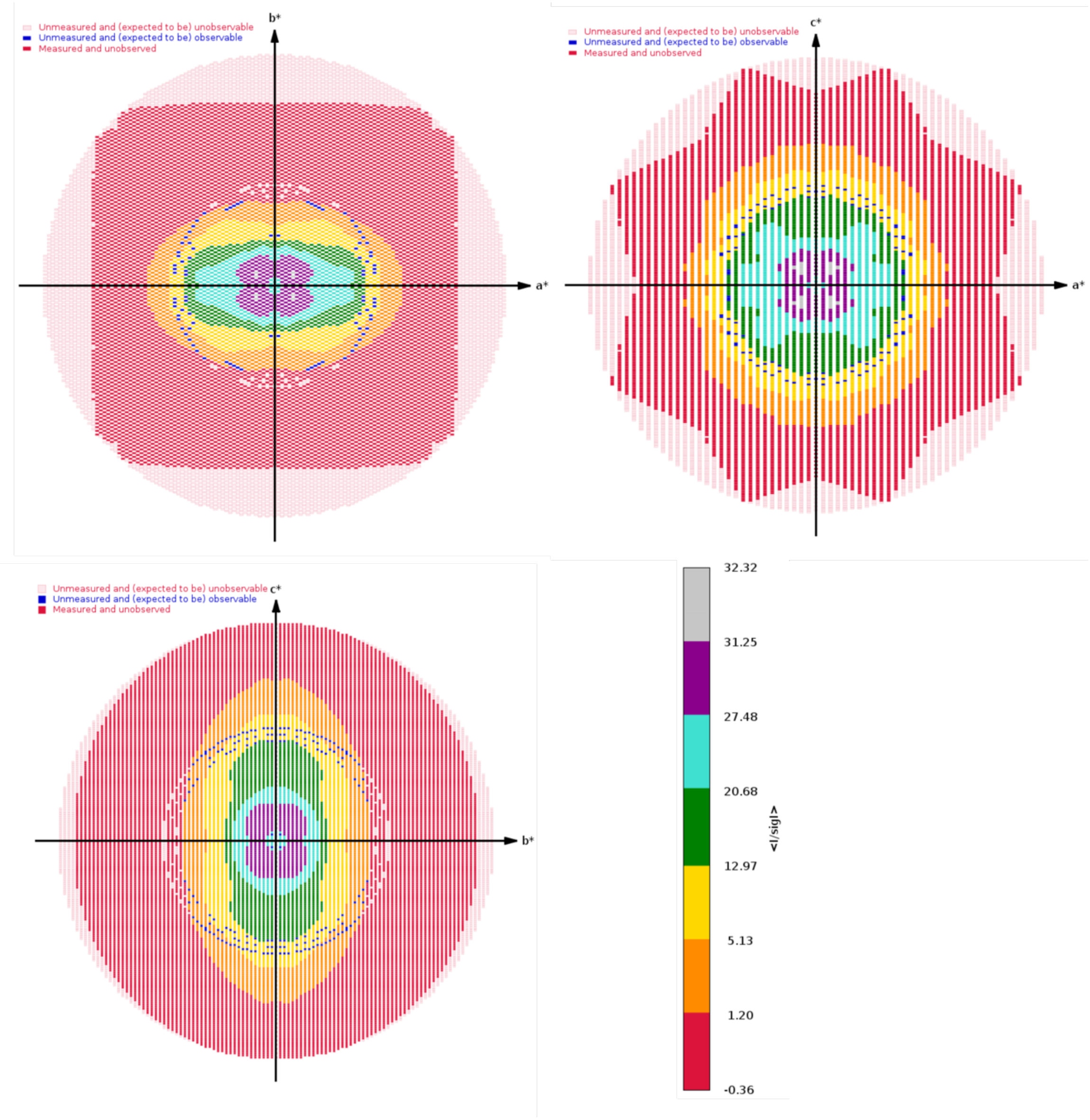


***Supplemental Figure 4*** *STARANISO calculated plots of reciprocal lattice points along the (a*,b*), (a*,c*) and (b*,c*) planes from the Δ80-PmHS2-**4-mer** dataset, colored by local mean I/σ(I). An I/σ(I) ≥ 1.2 threshold was used as the anisotropic cut-off, which defines the resolution limits reported in **Supplemental Table 1**.*

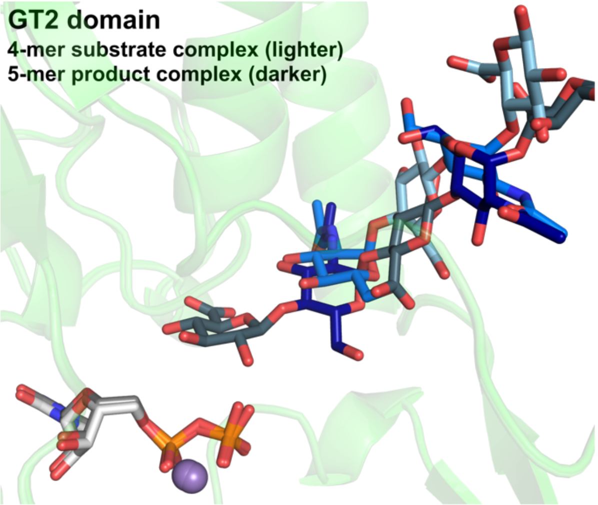


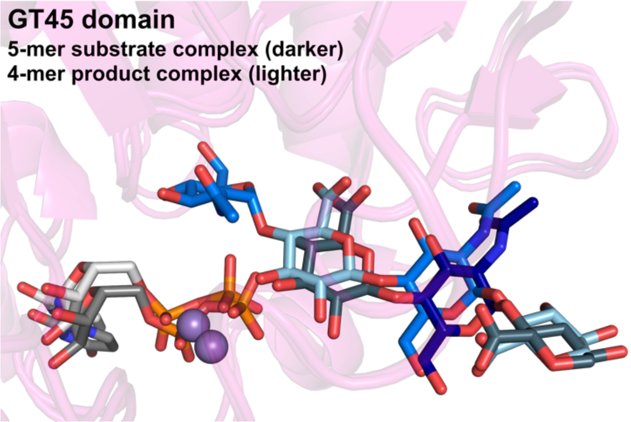


***Supplemental Figure 5*** *Overlays of* ***4-mer*** *and* ***5-mer*** *complexes within the GT2 and GT45 active sites. Ligands from the* ***5-mer*** *complex are colored darker. Minor shifts between substrate and product complexes can be observed, consistent with rearrangement of product ligand binding after enzymatic transfer.*

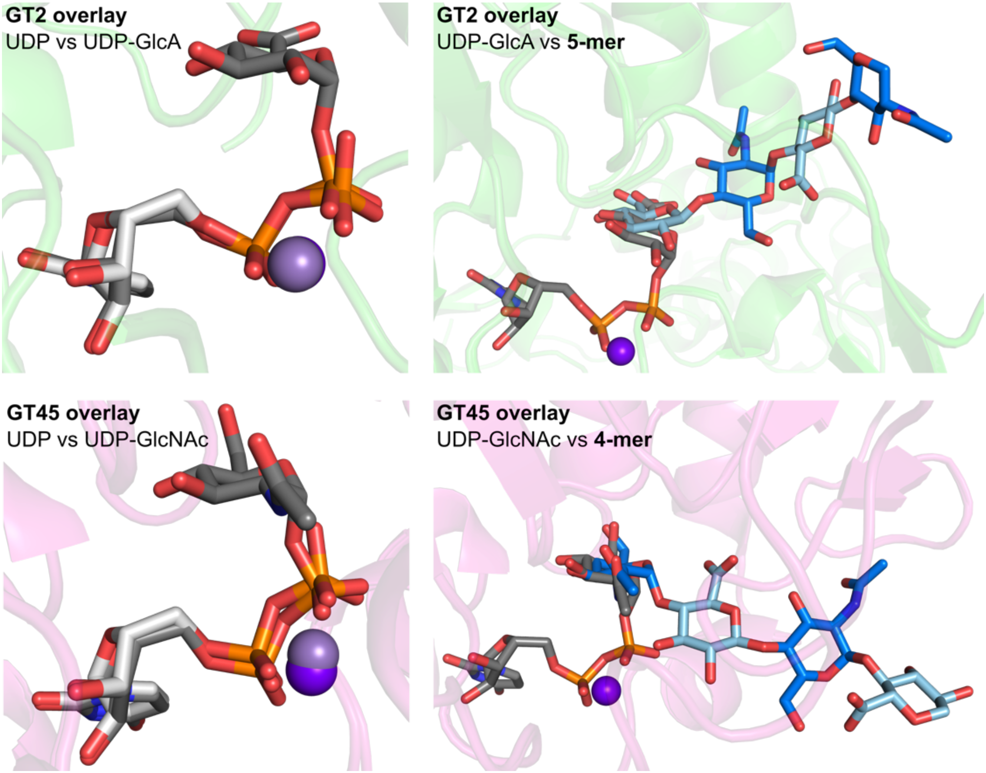


***Supplemental Figure 6*** *Comparison of UDP, UDP-sugar and oligosaccharide binding modes within the GT2 and GT45 active sites. UDP-sugars coordinate to* $Mn^{2+}$ *in an essentially identical fashion to free UDP. UDP-sugar complexes also display good sugar overlap with heparosan product ligands* ***5-mer*** *and* ***4-mer*** *within the GT2 and GT45 domains respectively.*

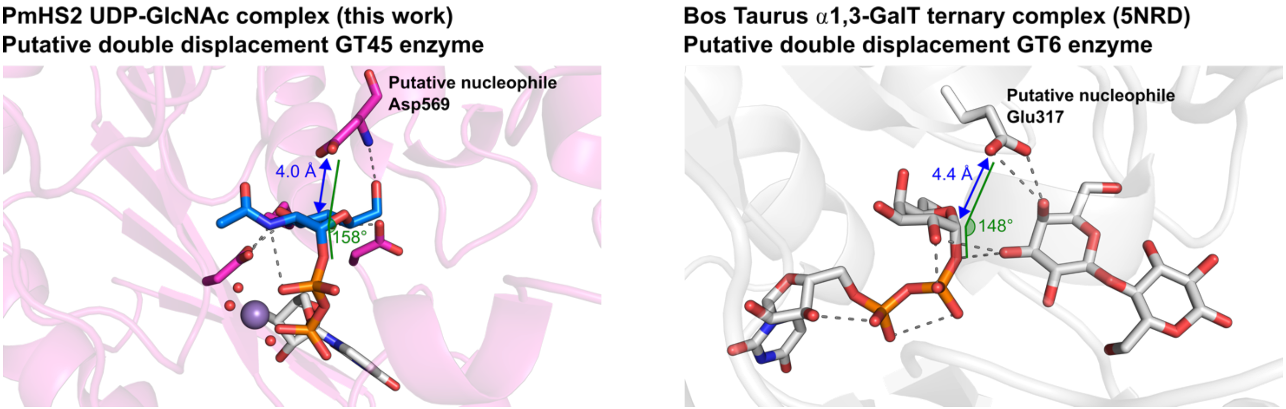


***Supplemental Figure 7*** *Comparison of putative nucleophile Asp569 in PmHS2 GT45 domain with putative nucleophile Glu317 in Bos Taurus α3-GalT, a member of the GT6 CAZy family. Asp569 shows more favorable distance to its UDP-sugar substrate for nucleophilic attack (4.0 Å vs 4.4 Å), as well as a more favorable angle of attack (158 ° vs 148 °).*

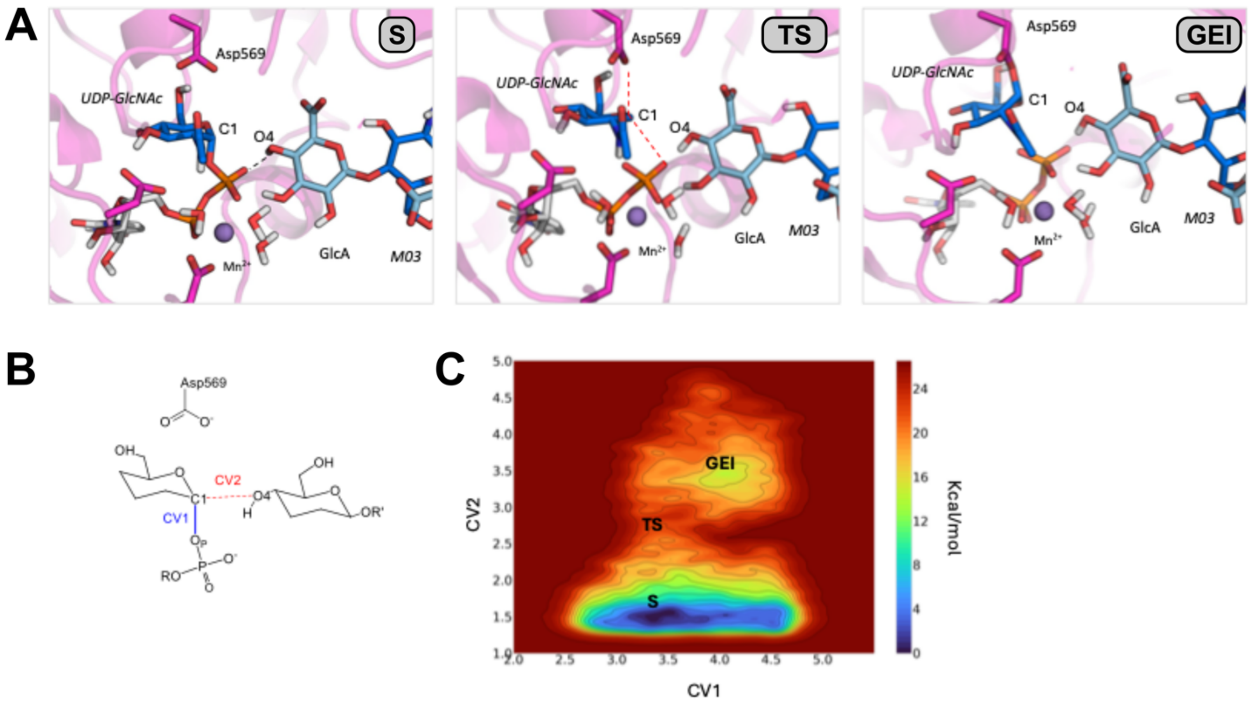


***Supplemental Figure 8*** *QM/MM simulation of GT45 reaction from* ***S****, using CVs optimized for* $S_Ni$*-like transfer. The initial structure was obtained from structural overlay of the enzyme complex with the UDP-GlcNAc donor and the enzyme complex with the M03 acceptor.*

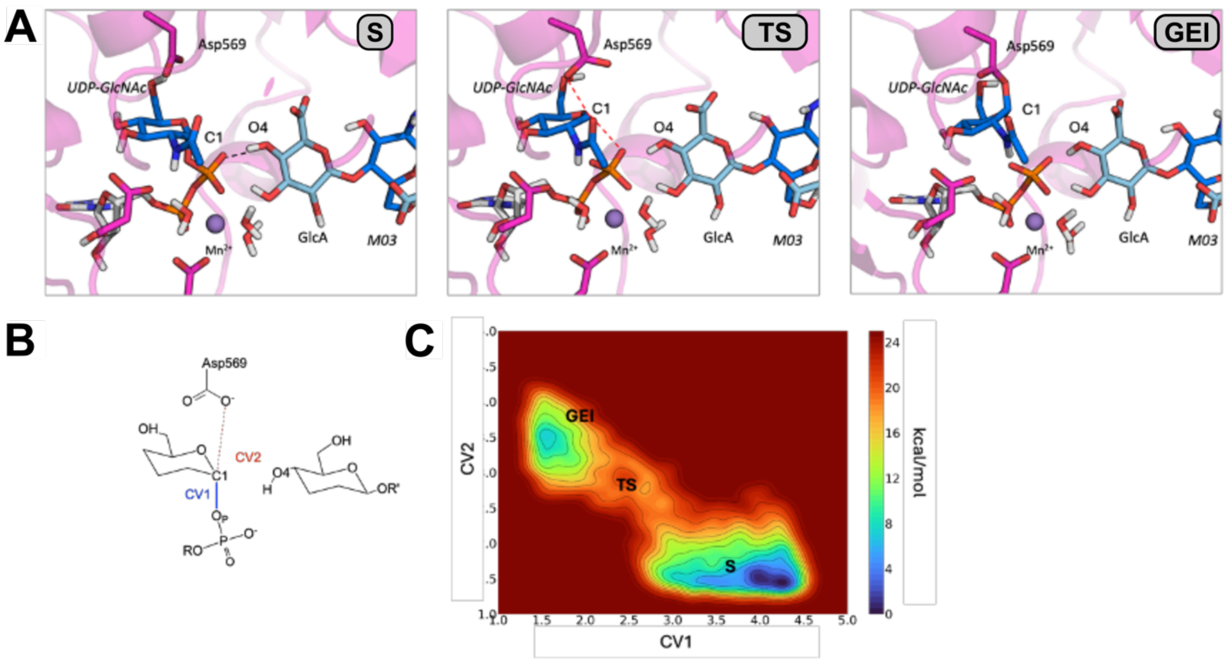


***Supplemental Figure 9*** *QM/MM simulation of GT45 reaction from* ***S****, using CVs optimized for double-displacement transfer.*

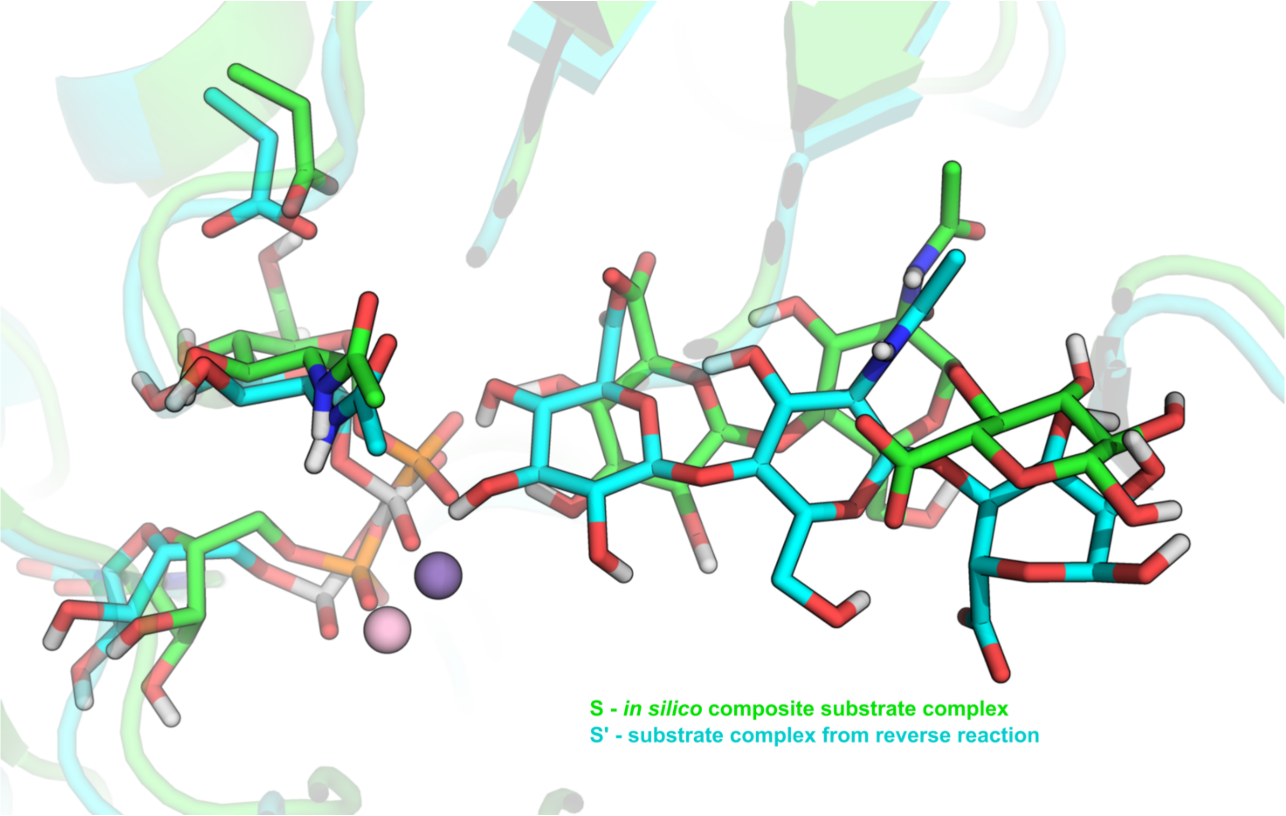


***Supplemental Figure 10*** *Structural overlay of ternary substrate complex* ***S*** *(green; in silico composite of UDP-sugar and* ***5-mer*** *oligosaccharide) against* ***S'*** *(cyan; substrate complex following reverse reaction from* ***P****). Differences in the pose of the oligosaccharide likely underscore the inability of QM/MM simulations to progress* ***S*** *towards a fully reacted product state.*

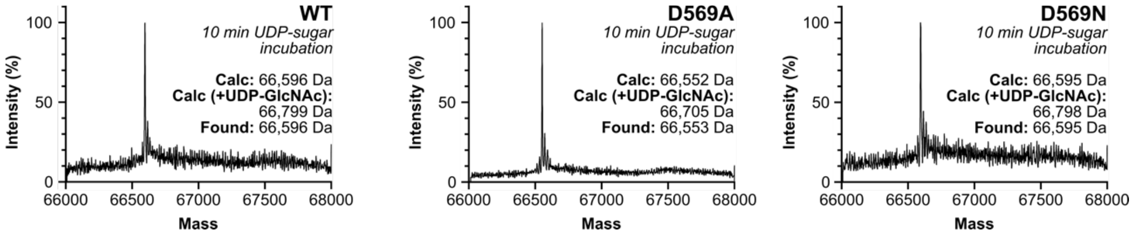


***Supplemental Figure 11*** *ESI-MS of wild-type Δ80-PmHS2 reaction with UDP-GlcNAc and UDP-GlcA. No additional peaks are observed with WT Δ80-PmHS2, Δ80-PmHS2(D569A), and Δ80-PmHS2(D569N).*

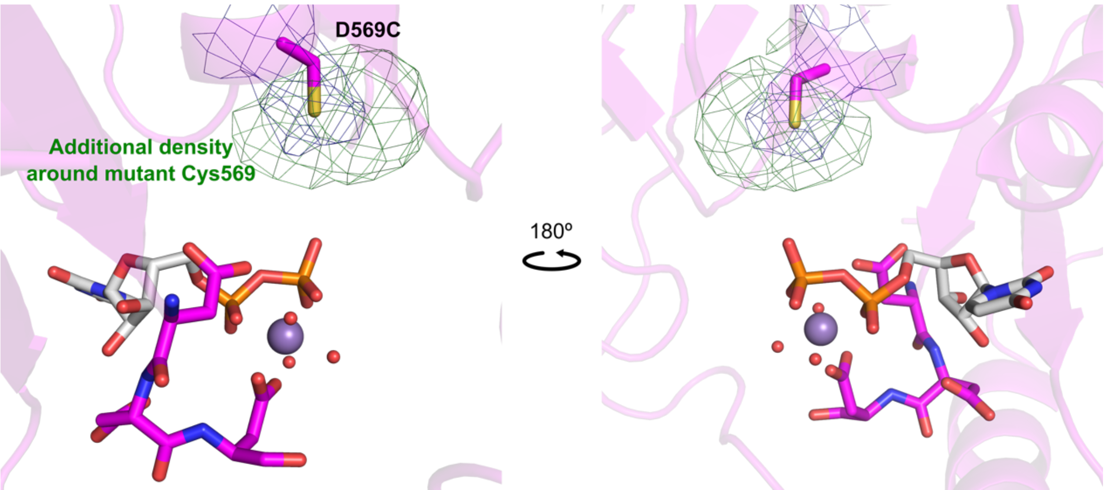


***Supplemental Figure 12*** *Δ80-PmHS2-D569C crystal structure shows additional difference density around the mutated Cys569 sidechain, consistent with the presence of a GlcNAc adduct. Electron density for the Cys569 sidechain is REFMAC $\sigma_A$-weighted 2mFo–DFc, contoured to 2.0 σ (0.17 $e^-$.$Å^{-3}$). Additional unmodeled electron density is REFMAC $\sigma_A$-weighted mFo–DFc, contoured to 2.5 σ (0.10 $e^-$.$Å^{-3}$).*

```
Prosthecochloris_sp._ARM31054.1                  shiiqftssrd-igdrrdharftdalky---gqghyyvcvdddivypadyierlagrset   127
Rhizobium_lusitanum_QND47458.1                   --kirhfidtd-gsrygdagkfwgleqv---e-davyvtcdddiiypddfvarmigelaq   352
Tritonibacter_mobilis_VCU59454.1                 --kikytisee-tanfrevgrfwglgqv---e-daifvtcsddiiyptdftarmvaelaq   351
Methylorubrum_populi_QDI82180.1                  --sinvfcsqd-yddigdagkfftfdps---r-nefyfsfdddivypndyadrlteaayh   296
Aeromonas_salmonicida_WHF42780.1                 --kityil--dtdsnkraaakffwvqst-----qgihltcdddifypnnyvknmvdelkk   205
Uncultured_marine_group_II/III_AIF03053.1        ---inavmsqneagdlgdrgkfywceqi-----rgyhltvdddilyppdyvgkilfnlrr   113
Sediminicoccus_rosea_UPY37480.1                  ---vefrrsqd-wddkgdagkfgwvdal---eepgyriivdddlvfaphftrvmveglrr   309
Methylomagnum_ishizawai_BBL74417.1               --kitlkfsqr----gypkakfamiesi---e-egiilsidddiaypedyaktlvaglrk   86
Salicibibacter_halophilus_QDI92864.1             ---kir-ltfs-nrnkggqmalrgvpqt-----sgyyfcvdddllypddyvdkmitfmkk   98
Shouchella_clausii_QNM41502.1                    ---kir-lsfs-tqnlggqmafkgfpet-----egyyfcvdddliypdnyvermvalmks   98
Halomonas_hydrothermalis_BCB08496.1              ---vlvissldtgvdmgdagkfyklssf---s-nvyyfscdddliypcnyvsemksflkk   441
Uncultured_virus_ASF00396.1                      ---ivth---i-gkdlnatgklfwalnp-----deyyfcidddirypstyaqdmiqylnt   97
Agrococcus_sp._UOV99866.1                        ---nv--rfft-gedlgdrakflflegf-----dgyyltcdddieyarfhvasiidgier   598
Fulvivirga_lutea_QSE98406.1                      ---vdifrsqe-hgdladngkfy-sppk-----ngfhltfdddilypsdyvktllsklkd   391
Viral_metagenome_1_QJA57629.1                    ----nvecell-dnslgdaarylhfged-----egysffcdddliyspnycrymihkynq   98
Viral_metagenome_4_QJH98368.1                    ----nvecell-dnslgdaarylhfged-----egysffcdddliyspnycrymihkynq   98
Viral_metagenome_2_QJA79022.1                    ----nvecell-dnslgdaarylhfged-----egysffcdddliyspnycrymihkynq   98
Gimesia_alba_QDT44047.1                          ---ielif--g-dnsrkgstkffwadkv-----qgyiltvdddliypsdyvevmcsaidk   591
Planctomycetes_bacterium_QDU09834.1              ---ielif--g-dnrrkgstkffwadki-----tgyiftcdddlvypsdyvqvmstaidr   465
Pseudomonas_inefficax_WNN38196.1                 ---itvih--d-kgdvgdrgkfyglinf-----egyyftcdddisyppyyvkhcidgier   701
Comamonas_testosteroni_WQG65134.1                ---ilveg-fg-kedlgdvgkffwvdqh-----dgiyfscdddliypkdyvdrtveklke   166
Sinorhizobium_meliloti_AEG56760.1                ---itvarsqe-hgelgdagkffwieqf-----dgfyfscdddivypayyverlvdklkd   567
Viral_metagenome_3_QJI01188.1                    ---ifinidmd-gidrgdldkfn-fdvl-----egytlycdddliyprnyvkemikaidq   98
Viral_metagenome_5_QJA94851.1                    ---ifinidmd-gidrgdldkfn-fdvl-----egytlycdddliyprnyvkemikaidq   98
Legionella_clemsonensis_ASQ47047.1               ---iqiftsqe-fgergdigkfyqvera-----fgyiftldddliypadyvstlvdkidt   99
Limnospira_fusiformis_ULB45240.1                 ---iyvarsqv-hgnlgdsgkffwceei-----qgyhftcdddlayppdyiktligkieq   1170
Freshwater_metagenome_CAB4147923.1               ------------nkdltdngkfygltlq---kkacyyfscdddiiyppnyiqhtiqaiek   289
Halovulum_dunhuangense_QIE55638.1                --nvtvldgrehrgdlsangkiyplkfv----rdsivltldddfifppdyvqtycdliak   110
Sulfitobacter_sp._UWR28334.1                     ---iravlgrdargdlgdtgrfydfpdq-----dayvftledslrypadyaarmiyhiem   445
Roseivivax_marinus_UMA67226.1                    --evtitr-sqdvgnlrdtgkflglrdv---sdstlfftidddiiypsnyvekmtaaldh   239
Ruegeria_sp._AXT28107.1                          --evtitr-sqdvgnlrdtgkflglrdv---sdstlfftidddiiypsnyvekmtaaldh   239
Parasedimentitalea_psychrophila_WIY26315.1       --kiicqf-gqqhgglkaegkflglald---ddaqiyvsadddmlypksfvrhlarmcil   101
Candidatus_Magnetococcus_CRH08300.1              --rvtvth-sadmpgyhangkllgvnqv---qqectylcidddiyfpydfaarmaqalsq   130
Methylovirgula_ligni_QAY96901.1                  --rvvpil-srdvpglqgngkflgltle---ksdclylgvdddiayprdyvrrtsealta   121
Paracoccus_aerodenitrificans_WBU63090.1          --kv---taiipnrdvkdagkfwfkp-d----pddivftidddigyppdyvaqtiaqfet   98
Pontivivens_ytuae_QPH52862.1                     --nl---npiipadntrdtgkfmpdv-e----dadlvflidddqiyppdyvdyarrsvek   76
Rhodovulum_sp._ARC87926.1                        --nv---nairpdedtkdtgkflpdy-r----gaewvflldddivfpehyvkdsiaamra   98
Dinoroseobacter_shibae_ABV94595.1                --nv---vpilidrdlkdtgkfwpsv-e----daewvvlidddlhypddyvattlarata   98
Actibacterium_sp._ALG89943.1                     --nv---tavipdedlkdvgkylpds-s----aaehvftmdddilypadyvattlsrlaa   98
Oceanicola_sp._QDC08797.1                        --ki---aqiipdedqkdvgkfypdv-s----gadwvfmidddmiyppdfvansiaamea   98
Pelagibacterium_sp._WMT91983.1                   --nv---ryvipdndlkdtgkfleprrs-----deyvftldddlkfppdyaatlidcyes   101
Alteromonas_macleodii_AFT79830.1                 --kl---eaiipdtdqrdvgkfldidvd----dedvilyvdddiiypenyalvmcqkl--   97
Escherichia_coli_QID72483.1                      --kl---npvipdkdykdvgkfifpcak-----ndmivltdddiiyppdyvekmlnfyns   98
Avibacterium_paragallinarum_QJE23390.1           --nl---npiipnkdykdvgkfihpivl-----dasyilvdddiiyppnyvehlqyfyek   98
Neisseria_sp._UO077238.1                         --kl---npiipktdykdvgkfihnvas-----ndeiiilidddiiypkdyvatlrdyyhq   98
Achromobacter_insolitus_AXA70137.1               -ikitvshsknfdidyrdnakfinfnklkekg-effyftcdddliypfdyvraliadiva   484
Moraxella_bovoculi_AKG08893.1                    -dkitvicsygksnslrdngkfilleelakngkngyyftidddidypldyvntlikkiny   464
Mergibacter_septicus_AWX13666.1                  -hkakvvicndkntslrdngkflllleelikenkdgyyftcdddieypvdyvntllerlyk   460
Pasteurella_multocida_AY292200.1                 -NKATVVHCKDKDNSIRDNGKFILLEELIEKNQDGYYITCDDDIIYPSDYINTMIKKLNE   498
Microvirga_lotononidis_WQ026097.1                ---it-alpasrledlrdigkfyhmsgv----ptgyfftvdddieypddycdtlvrkied   484
Pseudoxanthomonas_mexicana_UOV00684.1            ---vv-vk--iggqdlrdngkfyftpvv----pdgycftvdddivypkdyvqglirkiel   1022
                                                                                     .  .*.  :   .

Prosthecochloris_sp._ARM31054.1                  lvdvagigtlfcpaelieedf-ss---ppygfsdlvlakyfftrsipvvairrqkswlkg   225
Rhizobium_lusitanum_QND47458.1                   rvhvaatgtgafhsslikmtl-ad--frhrnmadiwlaeylhrhgipayavprkegwlls   449
Tritonibacter_mobilis_VCU59454.1                 rvhipattacafhtktvpvsf-ae--cdpgipadaqiatyaethsisryviprpaawlqt   448
Methylorubrum_populi_QDI82180.1                  rvhvlgtgtlcippgifkhkp-p---fyfrnmadiwiarqlskqgtgqicvarayewlkp   391
Aeromonas_salmonicida_WHF42780.1                 qvhlagtgtlcfrtktlqkke-fdnlwnfaastdewlacfsktnsiklilinrtknwmks   303
Uncultured_marine_group_II/III_AIF03053.1        evaligtgvlaydanelavdf-sk--fksnywadgwfslqarnanfrcivirhkkdwlkp   206
Sediminicoccus_rosea_UPY37480.1                  tvhvlgtnafcahsdtlklrw-ad--fghrnmadiyvallaqkrgipmvalerprhlawq   406
Methylomagnum_ishizawai_BBL74417.1               yvhavgsgiaawdaryckvka-eh--fpfsagadsqlavlgqknrwpfvllshtakwlvp   180
Salicibibacter_halophilus_QDI92864.1             rvmipgvgtacmhtksfsisp-md--fahknmrdavvackaakegisvvaidrgenwikk   196
Shouchella_clausii_QNM41502.1                    qvlipgvgtscmhsdsfaitp-nd--fpyqnmrdafiackaakegvpiiaikrpenwink   196
Halomonas_hydrothermalis_BCB08496.1              pvhfggtgvmafhtsfvnfnf-nf--frtpnmadvwvglyaqenkipvmilphhenwikh   537
Uncultured_virus_ASF00396.1                      rvhvigngvsvfntnnvkidy-tk--fkyhymddievslqlqeqnipaiavahdsnylgy   195
Agrococcus_sp._UOV99866.1                        gvhllgtgvcgfhtdtvrptm-de--fvhpnmadaflaihaqrrevpmvvlahkggeamp   695
Fulvivirga_lutea_QSE98406.1                      yvhiigtgtlgyhtsllnfdl-se--vdkthmidiffglhcqknsipliciehpeawmqe   488
Viral_metagenome_1_QJA57629.1                    evetggtgvmlintkdmkini-ed--flrpnmadiwvakfaheqnvkimviehnanivhy   193
Viral_metagenome_4_QJH98368.1                    evetggtgvmlintkdmkini-ed--flrpnmadiwvakfaheqnvkimviehnanivhy   193
Viral_metagenome_2_QJA79022.1                    evetggtgvmlintkdmkini-ed--flrpnmadiwvakfaheqnvkimviehnanivhy   193
Gimesia_alba_QDT44047.1                          fvdilgtgtigfhtddikmsm-ad--mdlpdmediaifklsynhsytkvvvahpegwfqs   685
Planctomycetes_bacterium_QDU09834.1              fvdilgtgaiafhtddikmsm-dd--mdmpdmediaifklsynhsytkvvvahpegwfqs   559
Pseudomonas_inefficax_WNN38196.1                 avhvlgtgctafhtstiqlry-ed--fktsnmadiyfaliggqaqsipfvvlkhdagealp   798
Comamonas_testosteroni_WQG65134.1                pvhilgtgcsafhtkflkikk-sd--flhpnmadiffsikgqeqkipfivlahekdeite   263
Sinorhizobium_meliloti_AEG56760.1                pvhilgtgcaafhtstmkvrl-sd--fpepnmadvffailgqqqrvpfivvqhdkgelie   664
Viral_metagenome_3_QJI01188.1                    iavipgtgvmgyhtdliqkid-fl--eqpknmadihmgifarkqgipviirkhkgtwikh   194
Viral_metagenome_5_QJA94851.1                    iavipgtgvmgyhtdliqkid-fl--eqpknmadihmgifarkqgipviirkhkgtwikh   194
Legionella_clemsonensis_ASQ47047.1               ladipgtgtlafhtdniklnh-ti--fltpnmsdiwlaiyakqnnipvisvsrpdlwikq   195
Limnospira_fusiformis_ULB45240.1                 fvhhlgtgamayhtstievvr-ne--fqhinmadiwfsllaqkykiprivikhqegwlqy   1265
Freshwater_metagenome_CAB4147923.1               qvdiagtgvsafktsyfnpidlin--sehkkmsdvifsleakkqnkkiqiiihgqewike   383
Halovulum_dunhuangense_QIE55638.1                lcslagsgtfgfdqrtlrfdp-ds--ffagvmvdlrlsmlareqglpiwvlprpegwlrh   205
Sulfitobacter_sp._UWR28334.1                     fvdvlgmgtaawhssalkpal-ed--fasegqsrlwfaaaaarkdvplfimpadtgwvre   539
Roseivivax_marinus_UMA67226.1                    pvsclgtgttafqvgtlrpki-sd--frthgmadlyfaeiaknndvplitiarrsdwlyd   333
Ruegeria_sp._AXT28107.1                          pvsclgtgttafqvgtlrpki-sd--frthgmadlyfaeiaknndvplitiarrsdwlyd   333
Parasedimentitalea_psychrophila_WIY26315.1       kvdvlatngtvhlvkdlkfdc-rn--wtyrnqvdlnfleeaskqdvglvttmrprswtrp   197
Candidatus_Magnetococcus_CRH08300.1              svdvigtgaaafstatlsfdl-eq--wpkvsmtdlgvaiaaakanipmlciarehpmvin   226
Methylovirgula_ligni_QAY96901.1                  hvdvlgtgtvmfdtemlrfdv-rt--wphvnmldlyfaleaatanlaliclprrreyllp   216
Paracoccus_aerodenitrificans_WBU63090.1          pialigtgtvcargssipdlk-vm--epyvgladiaychwlfkqqilpwalprkagwlpr   194
Pontivivens_ytuae_QPH52862.1                     rvsqlgtghliarpsmlppfd-ym--kdakrfvdirfarwchekkipmlclarpegylnd   177
Rhodovulum_sp._ARC87926.1                        vadelgtgvmiskpehlpdfe-sv--rsaqkfidvrmarwcfenditqvclprpagylra   211
Dinoroseobacter_shibae_ABV94595.1                rvdqlgtgtvvlksehmppfd-fm--etsqkfvdvrfalwafqqdlaciclprprewlgv   208
Actibacterium_sp._ALG89943.1                     vvdqlgsgvsvmrgcdfppfd-fm--kgsqgfvdvrlarwchehgilpvclpraagwlsa   204
Oceanicola_sp._QDC08797.1                        vvdqiatnaavmrgedfplhs-ym--sgsqkfvdvrlarwcfergitpvclpraagwlge   210
Pelagibacterium_sp._WMT91983.1                   lvnqlgtgcamtrgdllapme-ym--assqrfvdvrfarychengigmvcvarpdqwigd   197
Alteromonas_macleodii_AFT79830.1                 lvnqigtgtvaikaryrpshk-fm--dgsqkyvdvrfavhcfknripiiciaredewlke   193
Escherichia_coli_QID72483.1                      vvnqlgtgtvflkadqlpslk-ym--dgsqrfvdvrfsrymleneigmicvpreknwlre   194
Avibacterium_paragallinarum_QJE23390.1           vvnqlgtgtvflkghqlpsls-ym--egsqkyvdvrfskylyennisllcipreanwqke   194
Neisseria_sp._UO077238.1                         ivnqlgtgtvylkghqlpglk-ym--dcsqsfvdvrfskymiengigmvciprsaawmhe   194
Achromobacter_insolitus_AXA70137.1               lvnnlgtgtvafhssifqsiepak--wkmggmvdiyfanealarqipmlcvarhagwlle   580
Moraxella_bovoculi_AKG08893.1                    lvhllgtgttafridlfqdfklsn--fthtgmadvffaiecqkrkipqiaisrfdnwise   561
Mergibacter_septicus_AWX13666.1                  pvnilgtgtvafrvgifsrfclse--fkfpgmadiyfaieckknnilqicierhnnwlke   557
Pasteurella_multocida_AY292200.1                 AVNVLGTGTVSFRVSLFNQFSLSD--FTHSGMADIYFSLLCKKNNILQICISRPANWLTE   595
Microvirga_lotononidis_WQ026097.1                kvnligtgtmafhsstiaf-e-pg--slpkgmadigvaiaakannvdmiavtrpekwltp   579
Pseudoxanthomonas_mexicana_UOV00684.1            vvdqlgtgtlafhtslwrpsc-eg--ytetgmvdawvavdarkrgipliaverkkawlap   1117
                                                         .
```

***Supplemental Figure 13*** *Multiple sequence alignment (Clustal Ω[1]) of the CAZy GT45 family – one representative from each genus within the database as of June 2024. PmHS2 is in green with capital letters. Putative nucleophile Asp569 is near-universally conserved across the GT45 family (cyan), with only Sulfitobacter sp. lacking this residue. The Sulfitobacter sp. sequence also lacks a DXD motif, suggesting it may not be a true glycosyltransferase.*

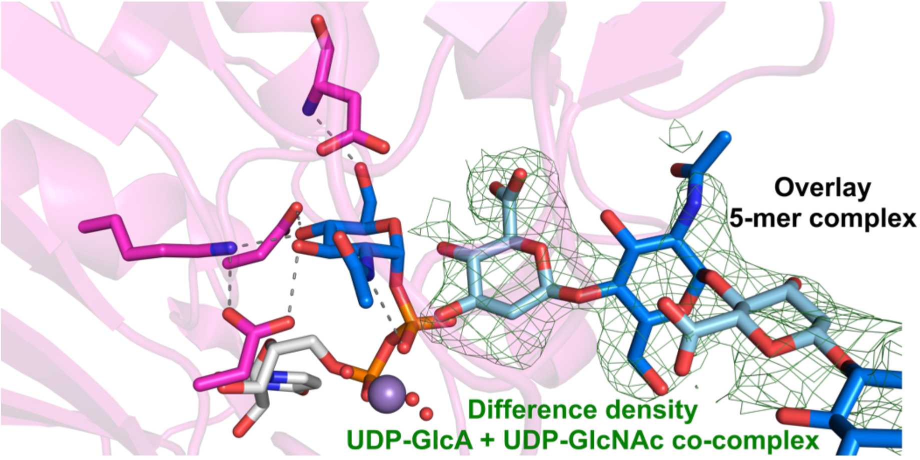


***Supplemental Figure 14*** *In crystallo polymerization of heparosan by Δ80-PmHS2. Incubation with UDP-GlcNAc and UDP-GlcA produces additional electron density within the GT45 domain, which matches the binding expected for a* ***5-mer*** *oligosaccharide.*

***Supplemental Table 1*** *Crystallographic processing and refinement statistics*

| | PmHS2- $Mn^{2+}$-UDP ***PDB*** 34ZL | PmHS2-$Mn^{2+}$-UDP-**4-mer** (STARANISO processed) ***PDB*** 34ZN | PmHS2-$Mn^{2+}$-UDP-**5-mer** ***PDB*** 34ZM | PmHS2-$Mn^{2+}$-UDPGlcNAc-UDPGlcA ***PDB*** 34ZO | PmHS2-D569C-$Mn^{2+}$-UDPGlcNAc-UDPGlcA ***PDB*** 34ZP |
|---|---|---|---|---|---|
| **Data collection** | | | | | |
| Space group | $C222_1$ | $C222_1$ | $P2_12_12_1$ | C121 | $C222_1$ |
| Cell dimensions | | | | | |
| *a*, *b*, *c* (Å) | 93.7, 191.3, 184.0 | 94.24, 187.43, 182.97 | 95.54, 182.03, 187.37 | 189.03, 94.45, 185.09 | 91.57, 186.81, 180.20 |
| α, β, γ (°) | 90, 90, 90 | 90, 90, 90 | 90, 90, 90 | 90, 99.29, 90 | 90, 90, 90 |
| Wavelength | 0.9762 Å | 0.9763 Å | 0.9763 Å | 0.9763 Å | 0. 9763 Å |
| Resolution (Å) | 46.88–2.10 (2.14–2.10) | 93.72–2.58, 4.02, 2.38* | 49.41–2.05 (2.09–2.05) | 59.80–2.16 (2.20–2.16) | 60.73–3.92 (3.99–3.92) |
| $R_{meas}$ | 0.20 (5.65) | 0.20 (1.89) | 0.17 (4.54) | 0.12 (5.92) | 0.95 (5.54) |
| $I / \sigma I$ | 6.1 (0.4) | 9.8 (1.6) | 11.5 (0.7) | 8.4 (0.3) | 3.2 (0.5) |
| Completeness (%) | 100 (100) | 90.0 (68.5) (ellipsoidal) | 100 (100) | 99.3 (98.5) | 100 (100) |
| Redundancy | 12.7 (11.5) | 13.8 (14.5) | 13.1 (13.5) | 6.6 (6.8) | 13.6 (14.4) |
| **Refinement** | | | | | |
| Resolution (Å) | 46.88–2.10 | 93.72–2.40 | 49.46–2.05 | 59.87–2.16 | 60.73–3.92 |
| No. reflections | 91459 | 30703 | 194157 | 162515 | 14242 |
| $R_{work}/R_{free}$ | 0.19/0.23 | 0.20/0.23 | 0.18/0.21 | 0.20/0.23 | 0.28/0.32 |
| No. atoms | | | | | |
| Protein | 9049 | 8998 | 17902 | 18043 | 8690 |
| Ligand/ion | 134 | 366 | 848 | 615 | 182 |
| Water | 332 | 98 | 1015 | 300 | 10 |
| *B*-factors | | | | | |
| Protein | 57.1 | 50.6 | 45.9 | 68.9 | 140.6 |
| Ligand/ion | 58.2 | 58.8 | 49.2 | 95.4 | 164.8 |
| Water | 50.7 | 34.1 | 46.0 | 57.3 | 151.3 |
| R.m.s deviations | | | | | |
| Bond lengths (Å) | 0.008 | 0.007 | 0.010 | 0.009 | 0.013 |
| Bond angles (°) | 1.47 | 1.56 | 1.58 | 1.70 | 1.16 |

*Resolution limits of anisotropic ellipsoid fitted to diffraction cut-off surface oriented along principal axes a*, b*, c*. See **Supplemental Figure 4**.

**Methods**

*Δ80-PmHS2 cloning*

A codon-optimized sequence for full length PmHS2 was ordered from IDT. From this, Δ80-PmHS2 was amplified using primers F – GATCGATCCATATGTTTGAAAAAATCTACACCTACAACC and R – GATCGATCGGATCCTTACAGGAAGTAGAACGGCAG, and subcloned into a pET15b backbone using the NdeI and BamHI restriction sites. The final construct encodes for a N-6xHis tag and thrombin cleavage sequence before the Δ80-PmHS2 gene. Asp569 mutants were generated by site-directed mutagenesis using a PCR based method[2]. All constructs were verified by Sanger sequencing before use.

*Protein production*

pET15b-Δ80-PmHS2 was transformed into BL21-Gold(DE3) cells (Agilent) for protein production. Cultures were grown in TB media supplemented with 100 μg/mL Ampicillin, induced using 0.5 mM isopropyl β-D-1-thiogalactopyranoside upon reaching an $OD_{600}$ of ~0.8, and further incubated at 16°C with shaking for 18 h. Cells were harvested by centrifugation at 4000 g, 4°C for 15 min, resuspended in Histrap buffer A (20 mM Tris pH 8.0, 500 mM NaCl, 20 mM Imidazole, 1 mM DTT) supplemented with cOmplete protease inhibitor (Roche) and DNase I (Sigma), and lysed using a CF1 cell disruptor (Constant Systems) at 35 kPSI. Lysate was cleared by centrifugation at 40,000 g, 4°C for 30 min, then directly applied to a 5 mL Histrap FF column (Cytiva) pre-equilibrated in Histrap buffer A. The loaded column was washed with 10 column volumes (CV) of Histrap buffer A, then eluted with Histrap buffer B (20 mM Tris pH 8.0, 500 mM NaCl, 500 mM Imidazole, 1 mM DTT) over a linear gradient of Histrap buffer A→B over 10 CV. Mutants were purified using the same process as for wild-type enzyme.

Δ80-PmHS2 containing fractions were pooled, and digested overnight at ambient temperature with 100:1 mass ratio Δ80-PmHS2:bovine plasma thrombin (Sigma). The digest reaction was rerun over a 5 mL Histrap FF column pre-equilibrated in Histrap buffer A, which was then washed with 4 CV Histrap buffer A. Flowthrough and wash fractions were pooled, concentrated to 2 mL using a Vivaspin 30 kDa molecular weight cutoff (MWCO) spin concentrator, then directly applied to a S75 16/600 column pre-equilibrated in SEC buffer (20 mM HEPES pH 7.4, 200 mM NaCl, 1 mM DTT). Purified Δ80-PmHS2 containing fractions were pooled and concentrated to 15–20 mg/mL using a Vivaspin 30 kDa MWCO spin concentrator. Aliquots of protein were flash frozen in LN2 for further use.

*PmHS2 enzyme activity assays – UDP-Glo*

The activity of 1 μM Δ80-PmHS2was determined in the presence of 500–0.16 μM UDP-GlcA and 500–0.16 μM UDP-GlcNAc, alone or in combination, in 25 mM HEPES pH 7.0, 150 mM NaCl, 5 mM $CaCl_2$, 5 mM $MnCl_2$. Reactions were incubated at ambient temperature in 96-well half area white plates (Greiner) for 320, 160, 80, 40, 20, 10 and 0 minutes, followed by the addition of UDP-Glo detection reagent (Promega), prepared according to the manufacturer's instructions. The reaction mixtures were incubated for a further 60 minutes, and luminescence measurements recorded using a CLARIOstar Plus microplate reader (BMG Labtech).

*PmHS2 enzyme activity assays – SDS-PAGE*

The activity of 1 μM Δ80-PmHS2 was determined in the presence of 1 mM UDP-GlcA and 1 mM UDP-GlcNAc, in 25 mM HEPES pH 7.0, 150 mM NaCl, 5 mM $CaCl_2$, 5 mM $MnCl_2$. Reactions were incubated at ambient temperature for 320, 160, 80, 40, 20 and 10 minutes, followed by heat inactivation at 95°C, and lyophilisation. Samples were analysed by PAGE, using 10%

polyacrylamide Tris-Acetate mini gels (8cm x 8cm) containing 17.5% (v/v) ethylene glycol, using Tris-MES running buffer. Subsequently, gels were treated with 0.1% (w/v) Cetrimonium bromide (Sigma) for 4 hours, followed by staining with 0.1% (w/v) Alcian Blue (Fisher) overnight. Gels were imaged using a ChemiDoc Imager (Bio-Rad) following extensive destaining in $H_2O$.

*SEC-MALLS*

SEC-MALLS experiments were run in SEC buffer. The injected sample comprised 100 µl Δ80-PmHS2 at 2.5 mg $ml^{-1}$. Experiments were conducted on a system comprising a Wyatt HELEOS II multi-angle light-scattering detector and a Wyatt rEX refractive-index detector linked to a Akta Pure system (Cytiva). Work was conducted at room temperature (20 ± 2°C). All solvents and buffers were 0.2 µm filtered before use and a further 0.1 µm filter was present in the flow path. The Unicorn software was used to control the Akta, and *ASTRA V* software was used for the HELEOS II and rEX detectors. All data were analysed using the *ASTRA V* software. Molecular masses were estimated using the Zimm fit method with a degree of 1. A value of 0.18 ml.$g^{-1}$ was used for the protein refractive-index increment (d*n*/dc).

*Intact protein mass spectrometry*

Δ80-PmHS2 samples were prepared for mass spectrometry analysis by direct dilution to 0.01 mg/mL protein in 0.1% formic acid, or desalting through Micro Bio-Spin 6 columns (BioRad) previously equilibrated in 0.1% formic acid.

Protein samples were analysed on Waters Xevo G2-XS QTof mass spectrometers equipped with a Waters Acquity UPLC. Separation was achieved using a Thermo Scientific ProSwift RP-2H monolithic column (4.6 mm × 50 mm) using water + 0.1% formic acid (solvent A) and acetonitrile + 0.1% formic acid (solvent B) as mobile phase at a flow rate of 0.4 ml/min and running a 5-min linear gradient of 5% solvent B to 95% solvent B for 5 min. Spectra were deconvoluted using MassLynx 4.1 (Waters) and the "MaxEnt1" deconvolution algorithm with the following settings: resolution: 0.1 Da per channel; damage model: uniform Gaussian; width at half height: 0.2 Da; minimum intensity ratios: 33% (left) and 33% (right); and iterate to convergence.

Timecourse reactions were set up with Δ80-PmHS2 wild-type or mutants, in reaction buffer comprising 25 mM HEPES pH 7.0, 150 mM NaCl, 5 mM $CaCl_2$, 5 mM $MnCl_2$. Reactions contained 1 mg/mL enzyme, and 10 mM each UDP-GlcNAc and UDP-GlcA. At defined timepoints, 10 µL of reaction mix were taken and processed as described above.

Δ80-*PmHS2-D569C In solution digestion*

Purified protein sample (20 µg) was resuspended in 100 mM ammonium bicarbonate, 10 mM TCEP, 2% sodium deoxycholate, pH 8, and incubated for 10 min at 95 °C, followed by centrifugation at 14,000 x*g* for 20 min at room temperature. Free cysteines were alkylated with 2-chloroacetamide (20 mM final concentration) in the dark at room temperature for 30 min. Protein sample was diluted with 100 mM ammonium bicarbonate to reduce sodium deoxycholate concentration to 1%. In-solution digestion was performed using chymotrypsin (Promega) at a protein-to-enzyme ratio of 50:1 (w/w) for 4 h at 37 °C. After the digestion, equal part of ethyl acetate was added to the sample, followed by acidification with formic acid to a final concentration of 2%. Sample was submitted to agitation for 1 min followed by centrifugation at 15,800 x*g* for 2 min. The organic phase was discarded, and the remaining aqueous phase was dried in speed vacuum concentrator. Peptides were resuspended in 0.1% formic acid and desalted using StageTip (Rappsilber et al., 2007) prior to LC-MS/MS analysis.

*Nano-Liquid-Chromatography, Mass Spectrometry and Data Processing*

Peptides were separated on Ultimate 3000 RSLCnano system (Thermo Fisher Scientific) equipped with a C-18 PepMap100 trap column (300 µm ID x 5 mm L, 100Å, Thermo Fisher Scientific) and an in-house packed Reprosil-Gold C-18 analytical column (50 µm ID x 500 mm L, 1.9 µm particle size, Dr. Maisch). Mobile phases (A: 0.1% FA, 5% DMSO and 94.9% water; B: 0.1% FA, 5% DMSO, 94.9% ACN) were delivered at a flow rate of 100nL/min. Separation was performed using a 15-min linear gradient (12-40% B). Eluting peptides were electro sprayed into an Orbitrap Ascend Tribrid mass spectrometer (Thermo Fisher Scientific) using data dependent acquisition (DDA) mode. Full MS scans (350-1,400 m/z) were acquired in the Orbitrap analyser at 60,000 resolution with a $1.2 \times 10^6$ AGC target, and 123 ms maximum injection time. The twenty most intense precursors (charge states 2-7) from MS1 scans were selected and isolated at 1.2 Th with the quadrupole for MS/MS event using higher-energy collision dissociation (HCD) at a normalized collision energy (NCE) of 26%, and the fragmentation spectra were detected in orbitrap at 7,500 resolution ($4 \times 10^4$ AGC target and 32 ms maximum injection time). Dynamic exclusion was enabled with the following settings: exclusion duration = 15 s, mass tolerance = ± 10 ppm, repeat count = 1.

LC-MS/MS raw data were processed using MaxQuant (version 2.3.0.0) against the Uniprot *Escherichia coli* protein database downloaded in September 2022 (4,372 sequences) plus the PmHS2 sequence. False-discovery rates were controlled at 1% both on peptide spectral match (PSM) and protein levels. Enzyme specificity was set to chymotrypsin. A maximum of two missed cleavages were allowed. HexNAc on cysteines (+203.08 Da), oxidation of methionine (+15.99 Da) and carbamidomethylation of cysteines (+57.02 Da) were set as variable modification.

*Protein crystallization and ligand complex generation*

Initial Δ80-PmHS2 crystallization trials were carried out using the PACT Premier (Molecular Dimensions), JCSG+ (Molecular Dimensions), Index (Hampton) and Morpheus (Molecular Dimensions) screens, testing unliganded protein, protein supplemented with 10 mM $MnSO_4$, or protein supplemented with 10 mM each $MnSO_4$ and UDP. Well diffracting crystals appeared in conditions containing Tris buffer and 2-Methyl-2,4-pentanediol (MPD), only in the presence of $Mn^{2+}$ and UDP. Final optimized crystallization conditions were 0.1 M Tris pH 7.8, 16% MPD, using 17 mg/mL Δ80-PmHS2 supplemented with 10 mM $MnSO_4$ and UDP. Crystals typically grew to size over 14 days.

Δ80-PmHS2-$Mn^{2+}$-UDP liganded crystals were harvested from crystallization drops and flash cooled in LN2 for data collection without further cryoprotection. **4-mer** and **5-mer** complexes were obtained by directly soaking ligand powder (Iduron) into drops containing Δ80-PmHS2-$Mn^{2+}$-UDP crystals, followed by 10–20 min incubation at ambient temperature, then harvesting and flash cooling in LN2 for data collection. UDP-sugar complexes were obtained by transferring crystals into fresh crystallization drops supplemented with 50 mM each UDP-GlcA and UDP-GlcNAc (Sigma), followed by 4–7 min incubation at ambient temperature, then harvesting and flash cooling in LN2 for data collection.

*Crystallographic data processing*

All diffraction data were collected at the Diamond Light Source (Harwell, UK). Data processing, model building and validation was carried out using programs from the CCP4i2 suite[3], unless otherwise specified. Diffraction images for the Δ80-PmHS2-$Mn^{2+}$-UDP, Δ80-PmHS2-$Mn^{2+}$-UDP-**5-**

**mer** and Δ80-PmHS2-$Mn^{2+}$-UDPGlcNAc-UDPGlcA complexes were processed using XDS[4] and AIMLESS[5]. Diffraction data for the Δ80-PmHS2-$Mn^{2+}$-UDP-**4-mer** complex displayed substantial anisotropy, and were processed using the STARANISO server[6]. The structure of the Δ80-PmHS2-$Mn^{2+}$-UDP complex was solved by molecular replacement using PHASER[7], applying a predicted structure from the ROBETTA server[8] as a search model. Other complexes were either solved by direct refinement against the Δ80-PmHS2-$Mn^{2+}$-UDP structure where isomorphous, or by molecular replacement using this structure. Solved structures were improved by iterative rounds of manual model building and refinement using COOT[9] and REFMAC5[10], respectively. Figures were prepared using Pymol.

*Sequence alignment and structure prediction.*
A list of GT45 sequences was obtained from CAZy, which consisted of 241 entries as of June 2024 (http://www.cazy.org/IMG/cazy_data/GT45.txt). To simplify analysis, only one sequence from each genus was used for alignment. 3 entries from the JGI (712030, 695, 2802) were discarded due to difficulties in finding the protein sequences. Two entries from Genbank (AUU83941.1, AUU83942.1) were also discarded, as they encoded for microproteins <100 amino acids that did not map to the catalytic regions of PmHS2. The final set for alignment contained 51 unique sequences, covering primarily bacterial genera. Multiple sequence alignments were carried out using the ClustalΩ[1] online interface. Structure predictions were carried out using the AlphaFold3[11] online interface.

*Reaction simulations – system preparation*
For the substrate complex simulation, an X-ray structure of Δ80-PmHS2 in complex with **5-mer** in each domain was taken as the initial structure, to which UDP-GlcNAc from the UDP-sugar cocomplex structure was overlaid. For the product complex simulation, an X-ray structure of Δ80-PmHS2 in complex with **4-mer** in each domain was taken as the initial structure. Terminal pNP groups were removed from every oligosaccharide, and the waters within the crystal structure were retained. The Δ80-PmHS2 homodimer was protonated at pH 5 with an internal dielectric of 10 using the H++ server[12,13]. The system was enclosed in a cubic water box spanning 10 Å away from the protein surface. The net charge was neutralized by adding two sodium ions. RESP charges of the UDP were calculated using Gaussian16 at the HF/6-31G* level of theory[14,15].

*Reaction simulations – classical MD simulations*
Classical MD simulations were carried out using AMBER2024, along with forcefields FF14SB[16,17], GLYCAM06[18] and TIP3P[19] for the enzyme, substrate, and water molecules, respectively. Input files were generated using AmberTools[20]. Time step was set to 2 fs for the production run, using the SHAKE algorithm[21]. The structure was minimized using Steepest Descent and equilibrated in several steps under NVT and NPT conditions. During NVT equilibration, the system was heated to 300 K using Berendsen thermostat[22] for 0.4 ns. During NPT equilibration, the system density was regulated for 0.4 ns using the Berendsen barostat. Electrostatic interactions were calculated using the Particle Mesh Ewald (PME) scheme, with an 8 Å cutoff.

*Reaction simulations – QM/MM MD*
QM/MM simulations were performed using CP2K v9.1[23]. A snapshot from the classical MD simulations was used as the initial structure. The QM region includes the Asp569, –1 and +1 sugar residues, phosphate group of the UDP, Manganese ion, and its immediate coordinators Asp481 and three water molecules. QM-MM boundaries include the C1-C2 bond of UDP-GlcNAc, C4-C5 and C2-C3 bonds of GlcNAc +2 sugar subsite, and the Cα -Cβ bonds of Asp residues. The residual

QM charge was redistributed over the MM atoms of the linked residues to retain a neutral charge in the MM region. The LJ parameters of H atoms were taken from the force-field GAFF2. The QM region was described by Density Functional Theory (DFT), using the Perdew-Burke-Ernzerhof (PBE) functional[24] along with molecular optimized Goedecker-Teter-Hutter (MOLOPT-GTH) pseudopotentials[25]. A triple-ζ valence polarized basis set functions (TZV2P) were used to expand the Kohn-Sham orbitals[26], with a 350 Ry cut-off. The conjugate gradients method[27] was used to optimize the structure, which was equilibrated under NVT conditions for 10 ps at 300K maintained by a thermostat using V-rescale algorithm[28] with a time constant of 10 fs.

*Reaction simulations – QM/MM metadynamics simulations*

The reaction mechanism was modelled with QM/MM metadynamics in two steps (**I** and **II**) both using a 2D CV space. The CVs are defined as the distances of two atomic pairs that a bond is assumed to be formed or broken: **I)** the bond between C1 and the glycosidic oxygen of the GlcA group ($d_{C1\text{-}O4'}$) and the distance between one O atom of the carboxylic acid group of Asp569 and the anomeric carbon ($d_{O\text{-}C1}$), **II)** the distance between one O atom of the carboxylic acid group of Asp569 and the anomeric carbon ($d_{O\text{-}C1}$) and the distance between O3 atom of the phosphate group and the glycosidic oxygen of the GlcNAc group (dC1-$O3_{(UDP)}$). CP2K, together with the Plumed 2.8 plug-in, was used to run the metadynamics. A simulation for **I** was launched with gaussian height of 1.0 kcal·mol-1, the gaussian width of 0.15 and 0.20 Å, and a deposition pace of 100 MD steps, corresponding to 50 fs. After successfully recrossing over the TS, a snapshot from the GEI was taken to conduct step **II**. In **II**, well-tempered metadynamics parameters for gaussian height was set to 2.0 $kcal.mol^{-1}$ and widths were set to 0.12 and 0.20 Å with a deposition pace of 100 MD steps and a bias factor of 30 at 300 K. Throughout all the metadynamics calculations, an upper boundary for distances of coordinating water molecules was set at 2.5 Å. FELs obtained from summation of hills were reweighted, discounting all the biases including metadynamics and restraints, using the same CVs for **I** and **II**, with a slight exception in one of the CVs of **II** by using the phosphorus atom of the phosphate group, instead of $O3_{UDP}$. Scott's rule was used to estimate the kernel bandwidths for the reweighting procedure.

*Free energy calculations*

For comparison of experimental *vs* calculated QM/MM free energy barrier, a GlcNAc-transfer $k_{cat}$ of 3.5 $s^{-1}$, taken from Na *et al* for Δ80-PmHS2[29], was used to inform transition state theory (TST) and the Eyring–Polanyi expression $k=(k_BT/h).exp[-G^{‡}(T)/RT]$, where k is the reaction rate, $k_B$ is the Boltzmann constant, h is the Planck constant, R is the gas constant. The formula assumes that the TST transmission coefficient of the reaction is 1, i.e. TS recrossing, tunneling and nonequilibrium contributions are neglected[30]. Because Δ80-PmHS2 is a homodimer, the $k_{cat}$ value used for the equation considered the reaction rate per site to be half the reported value *i.e.* 1.75 $s^{-1}$.

**Supplemental References**


1. Sievers, F.; Higgins, D. G., Clustal omega. *Curr Protoc Bioinformatics* **2014,** *48*, 3 13 1-3 13 16.

2. Liu, H.; Naismith, J. H., An efficient one-step site-directed deletion, insertion, single and multiple-site plasmid mutagenesis protocol. *BMC Biotechnol* **2008,** *8*, 91.

3. Potterton, L.; Agirre, J.; Ballard, C.; Cowtan, K.; Dodson, E.; Evans, P. R.; Jenkins, H. T.; Keegan, R.; Krissinel, E.; Stevenson, K.; Lebedev, A.; McNicholas, S. J.; Nicholls, R. A.; Noble, M.; Pannu, N. S.; Roth, C.; Sheldrick, G.; Skubak, P.; Turkenburg, J.; Uski, V.; von Delft, F.; Waterman, D.; Wilson, K.; Winn, M.; Wojdyr, M., CCP4i2: the new graphical user interface to the CCP4 program suite. *Acta Crystallogr D Struct Biol* **2018,** *74* (Pt 2), 68-84.

4. Kabsch, W., Xds. *Acta Crystallogr D Biol Crystallogr* **2010,** *66* (Pt 2), 125-32.

5. Evans, P. R.; Murshudov, G. N., How good are my data and what is the resolution? *Acta Crystallogr D Biol Crystallogr* **2013,** *69* (Pt 7), 1204-14.

6. Tickle, I. J., Flensburg, C., Keller, P., Paciorek, W., Sharff, A., Vonrhein, C., Bricogne, G. STARANISO (http://staraniso.globalphasing.org/cgi-bin/staraniso.cgi). Cambridge, United Kingdom: Global Phasing Ltd.

7. McCoy, A. J., Solving structures of protein complexes by molecular replacement with Phaser. *Acta Crystallogr D Biol Crystallogr* **2007,** *63* (Pt 1), 32-41.

8. Kim, D. E.; Chivian, D.; Baker, D., Protein structure prediction and analysis using the Robetta server. *Nucleic Acids Res* **2004,** *32* (Web Server issue), W526-31.

9. Emsley, P.; Lohkamp, B.; Scott, W. G.; Cowtan, K., Features and development of Coot. *Acta Crystallogr D Biol Crystallogr* **2010,** *66* (Pt 4), 486-501.

10. Murshudov, G. N.; Skubak, P.; Lebedev, A. A.; Pannu, N. S.; Steiner, R. A.; Nicholls, R. A.; Winn, M. D.; Long, F.; Vagin, A. A., REFMAC5 for the refinement of macromolecular crystal structures. *Acta Crystallogr D Biol Crystallogr* **2011,** *67* (Pt 4), 355-67.

11. Abramson, J.; Adler, J.; Dunger, J.; Evans, R.; Green, T.; Pritzel, A.; Ronneberger, O.; Willmore, L.; Ballard, A. J.; Bambrick, J.; Bodenstein, S. W.; Evans, D. A.; Hung, C. C.; O'Neill, M.; Reiman, D.; Tunyasuvunakool, K.; Wu, Z.; Zemgulyte, A.; Arvaniti, E.; Beattie, C.; Bertolli, O.; Bridgland, A.; Cherepanov, A.; Congreve, M.; Cowen-Rivers, A. I.; Cowie, A.; Figurnov, M.; Fuchs, F. B.; Gladman, H.; Jain, R.; Khan, Y. A.; Low, C. M. R.; Perlin, K.; Potapenko, A.; Savy, P.; Singh, S.; Stecula, A.; Thillaisundaram, A.; Tong, C.; Yakneen, S.; Zhong, E. D.; Zielinski, M.; Zidek, A.; Bapst, V.; Kohli, P.; Jaderberg, M.; Hassabis, D.; Jumper, J. M., Accurate structure prediction of biomolecular interactions with AlphaFold 3. *Nature* **2024,** *630* (8016), 493-500.

12. Gordon, J. C.; Myers, J. B.; Folta, T.; Shoja, V.; Heath, L. S.; Onufriev, A., H++: a server for estimating pKas and adding missing hydrogens to macromolecules. *Nucleic Acids Res* **2005,** *33* (Web Server issue), W368-71.

13. Anandakrishnan, R.; Aguilar, B.; Onufriev, A. V., H++ 3.0: automating pK prediction and the preparation of biomolecular structures for atomistic molecular modeling and simulations. *Nucleic Acids Res* **2012,** *40* (Web Server issue), W537-41.

14. Slater, J. C., A Simplification of the Hartree-Fock Method. *Physical Review* **1951,** *81* (3), 385-390.

15. Ditchfield, R.; Hehre, W. J.; Pople, J. A., Self-Consistent Molecular-Orbital Methods. IX. An Extended Gaussian-Type Basis for Molecular-Orbital Studies of Organic Molecules. *The Journal of Chemical Physics* **1971,** *54* (2), 724-728.

16. Maier, J. A.; Martinez, C.; Kasavajhala, K.; Wickstrom, L.; Hauser, K. E.; Simmerling, C., ff14SB: Improving the Accuracy of Protein Side Chain and Backbone Parameters from ff99SB. *J Chem Theory Comput* **2015,** *11* (8), 3696-713.

17. Tian, C.; Kasavajhala, K.; Belfon, K. A. A.; Raguette, L.; Huang, H.; Migues, A. N.; Bickel, J.; Wang, Y.; Pincay, J.; Wu, Q.; Simmerling, C., ff19SB: Amino-Acid-Specific Protein Backbone Parameters Trained against Quantum Mechanics Energy Surfaces in Solution. *Journal of Chemical Theory and Computation* **2020,** *16* (1), 528-552.

18. Kirschner, K. N.; Yongye, A. B.; Tschampel, S. M.; Gonzalez-Outeirino, J.; Daniels, C. R.; Foley, B. L.; Woods, R. J., GLYCAM06: a generalizable biomolecular force field. Carbohydrates. *J Comput Chem* **2008,** *29* (4), 622-55.
19. Jorgensen, W. L.; Chandrasekhar, J.; Madura, J. D.; Impey, R. W.; Klein, M. L., Comparison of simple potential functions for simulating liquid water. *The Journal of Chemical Physics* **1983,** *79* (2), 926-935.
20. Case, D. A.; Aktulga, H. M.; Belfon, K.; Cerutti, D. S.; Cisneros, G. A.; Cruzeiro, V. W. D.; Forouzesh, N.; Giese, T. J.; Gotz, A. W.; Gohlke, H.; Izadi, S.; Kasavajhala, K.; Kaymak, M. C.; King, E.; Kurtzman, T.; Lee, T. S.; Li, P.; Liu, J.; Luchko, T.; Luo, R.; Manathunga, M.; Machado, M. R.; Nguyen, H. M.; O'Hearn, K. A.; Onufriev, A. V.; Pan, F.; Pantano, S.; Qi, R.; Rahnamoun, A.; Risheh, A.; Schott-Verdugo, S.; Shajan, A.; Swails, J.; Wang, J.; Wei, H.; Wu, X.; Wu, Y.; Zhang, S.; Zhao, S.; Zhu, Q.; Cheatham, T. E., 3rd; Roe, D. R.; Roitberg, A.; Simmerling, C.; York, D. M.; Nagan, M. C.; Merz, K. M., Jr., AmberTools. *J Chem Inf Model* **2023,** *63* (20), 6183-6191.
21. Ciccotti, G.; Ryckaert, J. P., Molecular dynamics simulation of rigid molecules. *Computer Physics Reports* **1986,** *4* (6), 346-392.
22. Berendsen, H. J. C.; Postma, J. P. M.; van Gunsteren, W. F.; DiNola, A.; Haak, J. R., Molecular dynamics with coupling to an external bath. *The Journal of Chemical Physics* **1984,** *81* (8), 3684-3690.
23. Kuhne, T. D.; Iannuzzi, M.; Del Ben, M.; Rybkin, V. V.; Seewald, P.; Stein, F.; Laino, T.; Khaliullin, R. Z.; Schutt, O.; Schiffmann, F.; Golze, D.; Wilhelm, J.; Chulkov, S.; Bani-Hashemian, M. H.; Weber, V.; Borstnik, U.; Taillefumier, M.; Jakobovits, A. S.; Lazzaro, A.; Pabst, H.; Muller, T.; Schade, R.; Guidon, M.; Andermatt, S.; Holmberg, N.; Schenter, G. K.; Hehn, A.; Bussy, A.; Belleflamme, F.; Tabacchi, G.; Gloss, A.; Lass, M.; Bethune, I.; Mundy, C. J.; Plessl, C.; Watkins, M.; VandeVondele, J.; Krack, M.; Hutter, J., CP2K: An electronic structure and molecular dynamics software package - Quickstep: Efficient and accurate electronic structure calculations. *J Chem Phys* **2020,** *152* (19), 194103.
24. Perdew, J. P.; Burke, K.; Ernzerhof, M., Generalized Gradient Approximation Made Simple. *Phys Rev Lett* **1996,** *77* (18), 3865-3868.
25. Goedecker, S.; Teter, M.; Hutter, J., Separable dual-space Gaussian pseudopotentials. *Phys Rev B Condens Matter* **1996,** *54* (3), 1703-1710.
26. Kohn, W.; Sham, L. J., Self-Consistent Equations Including Exchange and Correlation Effects. *Physical Review* **1965,** *140* (4A), A1133-A1138.
27. Hestenes, M. R.; Stiefel, E., Methods of conjugate gradients for solving linear systems. *Journal of research of the National Bureau of Standards* **1952,** *49*, 409-435.
28. Bussi, G.; Donadio, D.; Parrinello, M., Canonical sampling through velocity rescaling. *The Journal of Chemical Physics* **2007,** *126* (1), 014101.
29. Na, L.; Yu, H.; McArthur, J. B.; Ghosh, T.; Asbell, T.; Chen, X., Engineer P. multocida Heparosan Synthase 2 (PmHS2) for Size-Controlled Synthesis of Longer Heparosan Oligosaccharides. *ACS Catal* **2020,** *10* (11), 6113-6118.
30. Garcia-Viloca, M.; Gao, J.; Karplus, M.; Truhlar, D. G., How enzymes work: analysis by modern rate theory and computer simulations. *Science* **2004,** *303* (5655), 186-95.
